# The reference-free pangenome of *Arabidopsis thaliana*

**DOI:** 10.1101/2025.06.25.661508

**Authors:** Lia Obinu, Andrea Guarracino, Urmi Trivedi, Andrea Porceddu

## Abstract

For several decades, scientists relied on unique reference genomes as the basis for various genomic studies. This methodology presents multiple drawbacks, particularly for variation studies, often yielding unrepresentative and incomplete observations regarding the diversity of the species or group under study. Pangenomes are collections of genomic sequences from several individuals of a species or population, which allow to overcome the limitations of studies based on a single reference genome.

In this study, we produced a 93-assembly pangenome of the model plant *Arabidopsis thaliana* using the reference-free method PanGenome Graph Builder (PGGB). The aim was to investigate the diversity within this species using this novel methodology, encompassing genomic sequences, genes, and pseudogenes. The pangenome exhibited a total length of 488.78 Mb, consisting of 30,391,243 nodes and 7,189,634 edges. We mapped a total of 2,325,577 genes and 23,894 pseudogenes across 93 assemblies, of which 36% and 0.9%, respectively, were classified as core. The observed variation in terms of genes and especially pseudogenes related to the geographical distance between sampling sites. Since the simulated pangenome growth curve based on gene did not reach a plateau, new accessions could potentially expand the represented diversity. This study enriches our understanding of intra-specific variation in *A. thaliana* and provides a new viewpoint on the potential applications of pangenomes in diversity research.

## 1 Introduction

For many years, researchers relied on single reference genomes as the foundation for diverse genomic investigations. This approach has numerous limitations, particularly as regards variation studies, frequently resulting in unrepresentative and partial results about the diversity of the species or group under study [1, 2].

The advancement of sequencing technologies, the reduction of costs, and the improvements of the assembly algorithms facilitated the production of many high-quality genome sequences in recent years, favouring the transition of genomics to the concept of “pangenome” [3, 4, 1, 5, 6]. Pangenomes are collections of genomic sequences of several individual belonging to a species or population. This concept, originally conceived for prokaryotes, has been extended to eukaryotes, including plants and animals [4, 7].

Pangenomes provide a comprehensive view of core sequences, present in all individuals, dispensable, present only in some individuals (with different levels of classification), and individual-specific sequences. This allows to capture gene presence/absence variation (PAV) and structural variants (SV) and polymorphisms, providing extensive knowledge about genomic diversity and insights into the relationship between genetic and functional diversity [8, 2, 9, 10]. They also enhance SNP discovery by providing a more complete reference, potentially improving genotype-phenotype associations [11].

Pangenomes can be used to investigate evolutionary history, population genetics, and phylogenetics. In plant genomics and agriculture, they are often used to identify genes associated with valuable traits such as disease resistance, yield components, and climate resilience and are particularly useful for breeding applications, genome editing, and biodiversity studies [2, 12].

*Arabidopsis thaliana* is a small flowering plant of the Brassicaceae family widely used as a model species in genomics mainly due to its small genome size, rapid life cycle, large number of offspring, easy cultivation, a selfing reproductive system, and a wide geographic distribution [13]. It was first sequenced in 2000 from the Columbia genotype, employing the minimum tiling path of BACs sequenced with Sanger technology [14]. The genome sequence spans approximately 135 Mbp in length and is organised into 5 chromosomes (2n = 10).

Several chromosome-level genome assemblies of this species were produced, especially in recent years (e.g., [15, 16, 17]), to complement the reference genome TAIR10 and to identify structural and functional differences. This makes this species a valuable resource to study pangenomes.

Previous works on *A. thaliana* pangenomes explored the evolutionary dynamics of the species [18], its local adaptations [19], and its intra-specific diversity [20], using 8, 32, and 73 genome assemblies, respectively. To obtain the gene-based pangenome, all these studies individually annotated each genomic sequence. To build the sequence-based pangenome, Jiao et al. (2020) [18], produced pairwise whole-genome sequence alignments of all possible pairs of genomes; Kang et al. (2023) [19], used Minigraph [21], which creates the pangenome in the form of a graph by setting one of the sequences as a reference-backbone and progressively aligning the others one by one to the reference; Lian et al. (2024) [20], performed a whole-genome alignment to the Col-PEK genome, chosen as a reference, using Minimap2 [22].

In this study, we generated a 93-assemblies pangenome of *A. thaliana* utilising the reference-free method PanGenome Graph Builder (PGGB), which constructs the pangenome as a variation graph through an all-to-all whole-genome alignment approach [23]. This strategy abandons reference-guided and tree-guided methodologies, delivering an unbiased representation of genetic variation. We utilised ODGI tools [24] to annotate the pangenome graph, rather than annotating each sequence separately, considering both genes and pseudogenes. Our objectives were to thoroughly investigate the diversity within this species utilising this novel methodology, encompassing not only genes and sequences but also pseudogenes, which was never attempted before; to evaluate the feasibility of directly annotating pangenome graphs; and to examine the correlation between the geographical distribution of the individuals considered in this study and their genetic diversity.

## 2 Materials and Methods

### 2.1 Data download and quality inspection

93 publicly available chromosome-level or complete genome assemblies of *A. thaliana* were downloaded from National Center for Biotechnology Information (NCBI) [25] and China National Center for Bioinformation (CNCB) [26] databases [16, 27, 18, 17, 28, 29, 30, 31]. The list of the assemblies, the collection sites of the samples, and their associated GenBank and project IDs are shown in Supplementary Materials Table S1. Among the assemblies, we included the reference genome TAIR10.1.

The assemblies were quality inspected using QUAST with parameters `--eukaryote, --large`, and `--k` [32]. Scaffolds smaller than 5 kbp and assembled organelles, when present, were excluded from the assemblies before performing any further analysis.

### 2.2 Pangenome graph building

The assemblies were renamed according to the PanSN-spec naming (https://github.com/pangenome/PanSN-spec) using Fastix (https://github.com/ekg/fastix). A multifasta containing all the assemblies was created. The multifasta was indexed using `samtools faidx` [33] and used as input for identifying the sequence communities with the PGGB function `partition-before-pggb` [23] with parameters -n 93, -p 90, and -s 10k.

Each community’s fasta file was used to build the pangenome graph using PGGB [23], with parameters -s 10000, -l 50000, -p 95, -n 93, -K 19, -F 0.001, -g 30, -k 23, -f 0, -B 10000000, -j 0, -e 0, -G 700,900,1100, -P 1,19,39,3,81,1, -O 0.001, -d 100, and -Q Consensus.

### 2.3 Sequence-based pangenome analysis

The characteristics of the pangenome graphs were obtained using the command odgi stats for each community [24]. A node coverage matrix for each graph was obtained using the command odgi paths with parameters -H, -D \#”, and -p1 [24]. The generated matrices were then processed with the in-house developed Python script nodes matrix processing.py (https://github.com/LiaOb21/arabidopsis_pangenome/blob/main/scripts/nodes_matrix_processing.py), which classifies nodes into genomic classes defined as: i. core, when they are found across all the 93 assemblies (parameter -na 93); ii. softcore, when they are present in at least 75 assemblies (parameter -sl 75); iii. dispensable, when they are found in between 2 and 74 assemblies; iv. and private, when they are found only in one assembly. The length of each node was obtained with odgi paths with parameters `--coverage-levels` 2,75,93 [24].

### 2.4 Pangenome reference-based annotation

Gene annotations for the reference genome TAIR10.1 were obtained from NCBI [25] in GFF3 format. From this file, only `gene` annotations were kept; therefore, CDS, exons, and other annotation types were excluded from the analysis. The annotations were split into subfiles based on the reference chromosome and saved in a BED format compatible with the PanSN-spec naming using Awk [34]. The BED subfiles contained the following columns: chromosome (in PanSN-spec format), start, end, and gene. Genes were recorded in the BED file as “gene name:start-end”.

Pseudogenes annotations were obtained from Mascagni et al. (2021) [35]. The file was first transformed into GFF3 format, as shown in `bed to gff3.ipynb` (https://github.com/LiaOb21/arabidopsis_pangenome/blob/main/scripts/bed_to_gff3.ipynb). For the pseudogenes annotated in the reverse strain, the forward coordinates were obtained for compatibility with the downstream analysis. The annotations were subsequently split into subfiles based on the reference chromosome and saved in BED format compatible with the PanSN-spec naming using Awk [34]. The BED subfiles contained the following columns: chromosome (in PanSN-spec format), start, end, and pseudogene. Pseudogenes were recorded in the BED file as “chromosome:start-end”, rather than using the names of the parent genes, i.e., the genes from where the pseudogenes originated.

The BED files were subsequently injected into the pangenome graph of the corresponding community using the function `odgi inject from ODGI` tools [24]. A step index for each graph was generated using `odgi stepindex` with parameters -a 0 [24]. The annotations were then mapped across the 93 assemblies using `odgi untangle` from the `untangle for annotation GitHub` branch [24]. These steps were performed separately for genes and pseudogenes.

The PAF files outputted by odgi untangle were processed using run collapse and merge.sh (https://github.com/LiaOb21/arabidopsis_pangenome/blob/main/scripts/run_collapse_and_merge.sh), a Bash wrapper that calls two Python scripts: collapse paf.py (https://github.com/LiaOb21/arabidopsis_pangenome/blob/main/scripts/collapse_paf.py) and merge paf.py (https://github.com/LiaOb21/arabidopsis_pangenome/blob/main/scripts/merge_paf.py). These two scripts were designed to join consecutive genomic regions annotated in the PAF file and to merge regions whose distance is lower than a given threshold (-d), respectively. run collapse and merge.sh processes each assembly separately and allows parallel execution of jobs to speed up the analysis. The script was run with parameters -d 100 and -j 20.

The processed PAF files obtained for the gene and pseudogenes annotations were analysed with core dispensable genes.py (https://github.com/LiaOb21/arabidopsis_pangenome/blob/main/scripts/core_dispensable_genes.py) and core dispensable pseudogenes.py (https://github.com/LiaOb21/arabidopsis_pangenome/blob/main/scripts/core_dispensable_pseudogenes.py) respectively, with parameters -cov 95 (coverage threshold), -id 95 (identity threshold), -jc 95 (Jaccard index threshold), -na 93 (number of assemblies), and -sl 75 (softcore limit). These two scripts classify genes and pseudogenes into genomic classes: i. core, when they are found across all the 93 assemblies; ii. Softcore, when they are present in at least 75 assemblies (-sl); iii. Dispensable, when they are found in between 2 and 74 assemblies; iv. and private, when they are found only in one assembly. The scripts output copy number variation matrices based on the results of the pangenome reference-based annotation.

As regards pseudogenes, the metrics were calculated based on the parent gene name, meaning that two pseudogenes matching the same functional locus were considered as being originated by duplications of that locus even if they contained different mutations.

The coverage and identity thresholds were chosen after several trials using different values, employing the community corresponding to chromosome 1 as example and considering genes annotations (Supplementary Materials Figure S1).

### 2.5 Non-reference sequences annotation

In order to obtain annotations for those sequences not covered by the reference genome, non-reference sequences were extracted from the pangenome graphs using `odgi paths` with the parameter `--non-reference-ranges` [24]. The output is a BED file containing the coordinates for all the regions in the non-reference assemblies that are not shared with the reference assembly. Using the script `filter private bed.py` (https://github.com/LiaOb21/arabidopsis_pangenome/blob/main/scripts/filter_private_bed.py) with parameter `-thr` 500 we separated the extracted regions into shorter than 500 bp and longer than 500 bp. The sequences shorter than 500 bp were enlarged using `bedtools slop` with parameter -b 500 [36] and joined again with the sequences longer than 500 bp. The resulting file was then processed using `bedtools merge` with parameter -d 100 to exclude overlapping and join close sequences [36]. The multifasta file corresponding to the final BED file was then extracted using `bedtools getfasta` [36] from the initial multifasta used to construct the pangenome graphs.

To identify potential coding sequences, the non-reference sequences were aligned against the UniRef 100 database [37] using `diamond blastx` with parameters `--range-culling, --top 0`, and -F 15 [38]. The alignments identified were subsequently processed with the script split and extract for exonerate.sh (https://github.com/LiaOb21/arabidopsis_pangenome/blob/main/scripts/split_and_extract_for_exonerate.sh), which divides diamond blastx results in chunks, extracts unique sequences and protein IDs, and extracts the corresponding FASTA sequences using seqtk subseq (https://github.com/lh3/seqtk). Each chunk was then analysed with Exonerate [39] launched through the Perl script PW exonerate.pl (https://github.com/LiaOb21/arabidopsis_pangenome/blob/main/scripts/PW_exonerate.pl), which runs Exonerate with parameters `--model protein2genome` and `--maxintron` 5000 on the alignments already identified by `diamond blastx` [38].

Exonerate results were then parsed using the script `parse exonerate.py` (https://github.com/LiaOb21/arabidopsis_pangenome/blob/main/scripts/parse_exonerate.py). This script considers as valid alignments only those showing at least 20% of coverage and 50% of identity. It classifies the alignments as genes, when they show 100% coverage and do not contain stop codons, or as pseudogenes, when they show coverage lower than 100% or possess stop codons. Pseudogenes that fall on the edge of the alignments are classified as “unclassified”, as we cannot know with certainty if they are present as complete genes in the assemblies. The results are stored in two different files, one containing the results for the UniRef100 IDs with a corresponding TAIR10 annotation and the other containing the results for the UniRef100 IDs with no corresponding TAIR10 annotation. We define as new genes and pseudogenes those that were annotated on the non-reference sequences and do not have a corresponding annotation on TAIR10.

The results for those UniRef100 IDs that did not have a unique corresponding TAIR10 annotation were excluded, as they could lead to misinterpretation of the results.

The unique UniRef100 IDs corresponding to new genes and pseudogenes were extracted and uploaded to UniProt ID mapping (https://www.uniprot.org/id-mapping), where a mapping UniRef100 to UniProtKB/Swiss-Prot was performed with the goal to obtain the IDs of reviewed proteins only, plus additional information. The results from the mapping were used to review the results obtained from the previous analysis, using the script review exonerate results.py (https://github.com/LiaOb21/arabidopsis_pangenome/blob/main/scripts/review_exonerate_results.py), which keeps only the results of reviewed proteins, i.e., those that mapped against UniProtKB/Swiss-Prot.

### 2.6 Final annotation screening

The Python scripts `final genes screening.py` (https://github.com/LiaOb21/arabidopsis_pangenome/blob/main/scripts/final_genes_screening.py) and `final pseudogenes screening.py` (https://github.com/LiaOb21/arabidopsis_pangenome/blob/main/scripts/final_pseudogenes_screening.py) were used to join the results coming from `odgi untangle` [24] and Exonerate [39] for genes and pseudogenes, respectively, with parameters -a 93 and -s 75. These scripts label genes and pseudogenes as core, softcore, dispensable, and private, as described above for the pangenome reference-based annotation. The scripts output the final statistics about how many genes and pseudogenes are present in each genomic class, and the final presence-absence matrices.

With this parsing, we identified new genes and pseudogenes, defined as features annotated on the non-reference sequences with UniRef100 IDs that do not have a corresponding TAIR10 annotation according to the official conversion table (https://www.arabidopsis.org/download/file?path=Proteins/Id_conversions/TAIR2UniprotMapping.txt). Furthermore, translocations were identified, meaning that genes annotated on a particular chromosome in TAIR10 were annotated on other chromosomes in other assemblies.

Figure 1 illustrates the main steps for the pangenome building and annotation.

**Figure 1:**
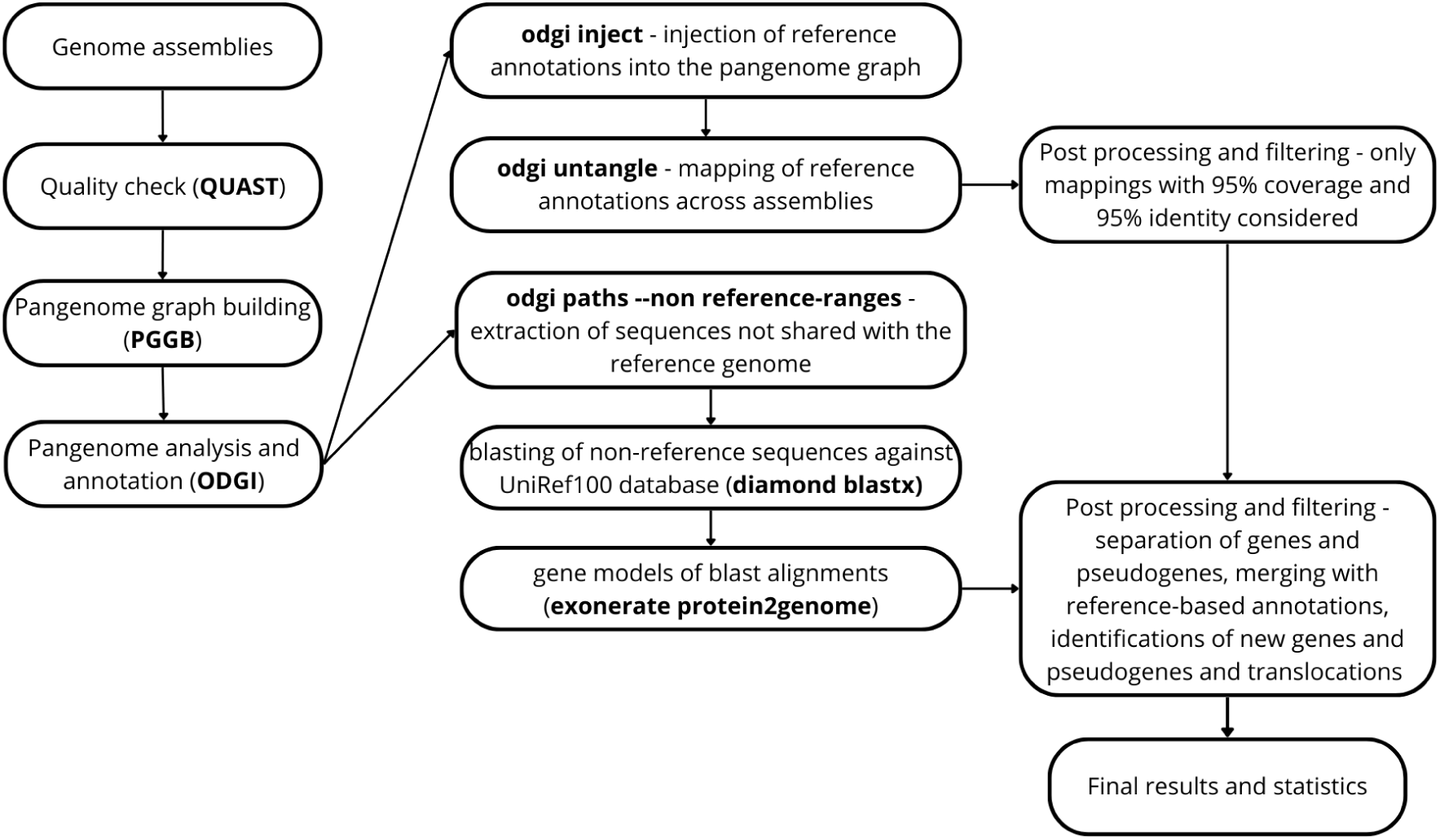
Flowchart of the main steps for pangenome graph building and annotation.

Considering the final matrix obtained for the genes, we simulated the growth of the pangenome as the number of accessions increased using the script `shuffle count.pl` (https://github.com/LiaOb21/arabidopsis_pangenome/blob/main/scripts/shuffle_count.pl).

### 2.7 Gene ontology enrichment analysis

We analysed gene ontology distributions among genomic classes taking into account core and variable (i.e., softcore, dispensable, and private simultaneously) genes and pseudogenes using TopGO [40]. We downloaded the Gene Universe of *A. thaliana* from https://www.arabidopsis.org/download/list?dir=GO_and_PO_Annotations. To study the enrichment of genomic compartments, we employed the Fisher exact test based on gene counts and the parent-child algorithms to deal with the GO structure.

### 2.8 Similarity analysis

The presence-absence matrices obtained from the final screening for each community were joined with the Python script `combine notransp matrices.py` (https://github.com/LiaOb21/arabidopsis_pangenome/blob/main/scripts/plot_scripts/combine_notransp_matrices.py) to obtain a single matrix for genes and pseudogenes. Jaccard distance matrices were obtained with the function vegdist from the R vegan package [41]. The hierarchical clustering was obtained with the function hclust from R stats package using the method `ward.D2` [42]. The hierarchical clustering was converted into a tree object with the as.phylo function from the R package ape [43].

The pangenome graphs were joined using the function `odgi squeeze` from ODGI tools [24]. A comprehensive distance matrix was then obtained using the function `odgi similarity` with parameters -D ‘#’, and -d. The Jaccard distances obtained were used to perform the hierarchical clustering with the function `hclust` from the R stats package using the default method (i.e., complete) [42]. The hierarchical clustering was converted into a tree object with the `as.phylo` function from the R package ape [43].

To evaluate the robustness of the trees, we calculated the cophenetic correlation, which examines the integrity of the dendrogram in preserving the pairwise distances from the original distance matrix. A cophenetic distance matrix was generated with the `cophenetic` function from the R stats package [42], and the correlation between the original distance matrix and the cophenetic matrix was calculated using the Pearson correlation coefficient using the ‘cor’ function from R stats [42].

To evaluate the relationship between genetic dissimilarities in terms of sequences, genes, and pseudogenes, and geographical distances, we conducted a Mantel test using the ade4 package in R [44, 45, 46, 47] with parameter `nrepet` = 9999.

### 2.9 Data visualisation

The statistics for genes, pseudogenes, and nodes categorised by genomic classes were visualised using Matplotlib [48].

The copy number variation of genes and pseudogenes for each assembly categorised by genomic classes was determined using the Perl script `comp node path CNV.pl` (https://github.com/LiaOb21/arabidopsis_pangenome/blob/main/scripts/plot_scripts/comp_node_path_CNV.pl) and visualised with the R package ggplot2 [49].

The number of genes, pseudogenes, and nodes considering their presence/absence for each assembly, categorised by genomic classes, was determined using the Perl script `comp node path PAV.pl` (https://github.com/LiaOb21/arabidopsis_pangenome/blob/main/scripts/plot_scripts/comp_node_path_PAV.pl) and visualised with the R package ggplot2 [49].

Similarity trees were visualised using the R packages ggtree [50, 51, 52, 53] and ggplot2 [49]. The tips of the trees were colour-coded according to the country of origin for each assembly sample.

The simulated pangenome growth curve was visualised with the scripts `curve plot.py` (https://github.com/LiaOb21/arabidopsis_pangenome/blob/main/scripts/curve_plot.py) and curve plot2.py (https://github.com/LiaOb21/arabidopsis_pangenome/blob/main/scripts/curve_plot2.py).

## 3 Results

### 3.1 Assembly quality assessment

The results of the quality assessment performed with QUAST are shown in Supplementary Materials Table S2. Overall, the assemblies showed a genome fraction, i.e., the percentage of aligned bases to the reference genome, between 85.35% and 99.99%, N50 between 22.59 Mbp and 30.45 Mbp, L50 equal to 3, and L90 equal to 5. Most of the assemblies contained 5 scaffolds, with the exception of those shown in Table 1, which were subjected to the removal of the assembled organelles or the filtering of scaffolds *<* 5 Kbp.

**Table 1:** Assemblies that were subjected to organelle removal or filtering of contigs smaller than 5000 bp before proceeding with the analysis.

| Assembly | Total number of contigs | Number of contigs after filtering | Initial length (bp) | Length after filtering (bp) | Applied filter |
| --- | --- | --- | --- | --- | --- |
| GCA_000001735 | 7 | 5 | 119668634 | 119668634 | Organelle removal |
| GCA_023115395 | 7 | 5 | 132081078 | 132081078 | Organelle removal |
| GCA_900660825 | 109 | 78 | 119626746 | 119530365 | Contigs <5000 bp |
| GCA_902460265 | 111 | 83 | 120129838 | 120059751 | Contigs <5000 bp |
| GCA_902460275 | 102 | 71 | 119749512 | 119657772 | Contigs <5000 bp |
| GCA_902460285 | 105 | 75 | 120338059 | 120246237 | Contigs <5000 bp |
| GCA_902460295 | 94 | 74 | 120289901 | 120224609 | Contigs <5000 bp |
| GCA_902460305 | 184 | 155 | 122202079 | 122112347 | Contigs <5000 bp |
| GCA_902460315 | 142 | 115 | 120795191 | 120728700 | Contigs <5000 bp |
| GCA_904420315 | 7 | 5 | 130173334 | 130173334 | Organelle removal |

### 3.2 Sequence-based pangenome

The pangenome consisted of five communities representing the five *A. thaliana* chromosomes. With a total length of 488.78 Mb, it included 1130 paths, 30,391,243 nodes and 7,189,634 edges (Table 2). Figure 2 represents the relative node composition of the *A. thaliana* pangenome. We classified 13% of nodes (3,939,502) as core, covering 78,057,221 bp (i.e., 16% of the total pangenome length); 22.2% (6,732,928) as softcore, accounting for 127,128,914 bp (26% of the total length); 46.9% (14,258,554) as dispensable, representing 22,032,786 bp (4.5% of the total length); and 18% (5,460,259) as private, resulting in a total length of 261,557,414 bp (53.5% of the total length).

**Figure 2:**
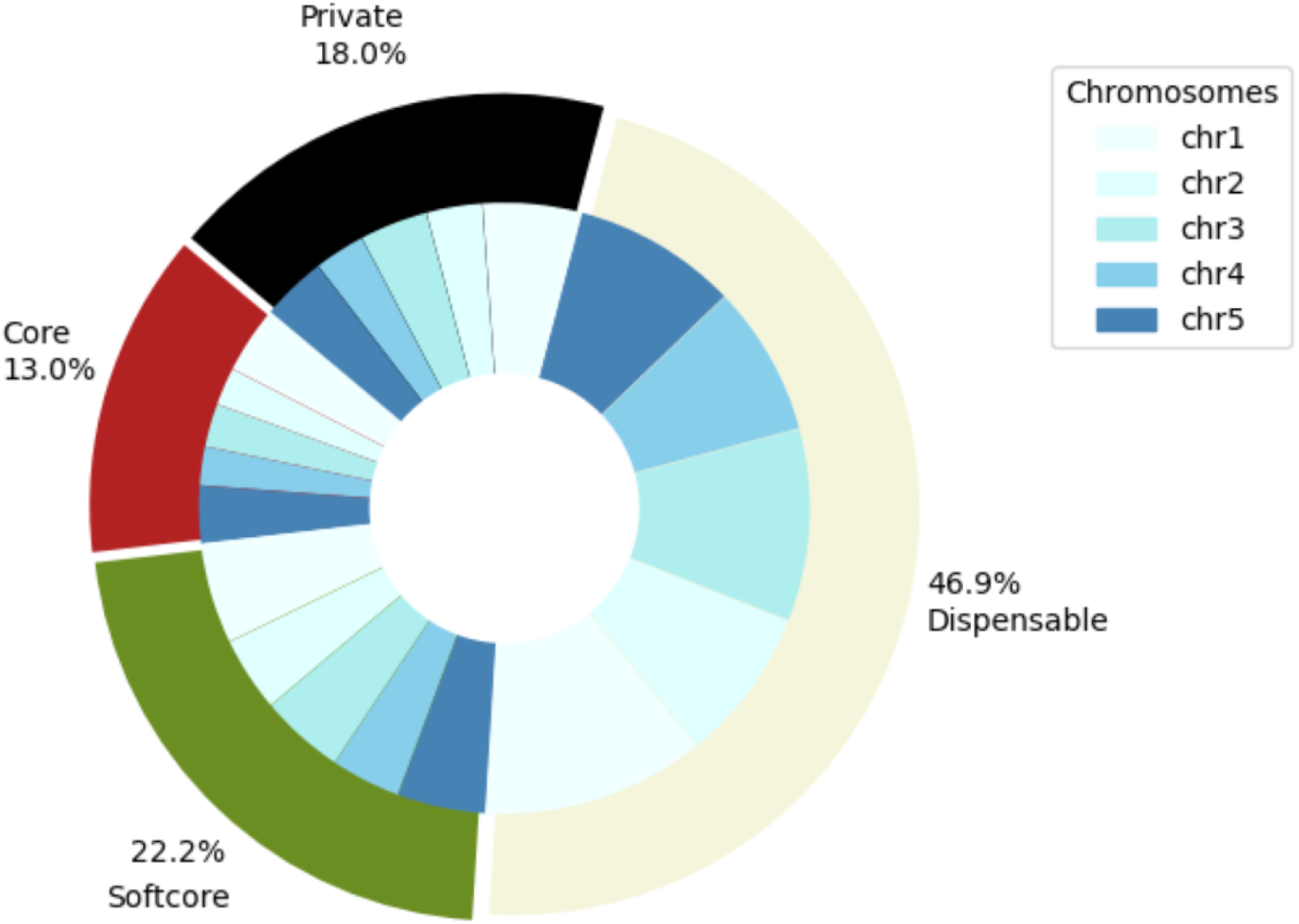
Relative node composition of the A. thaliana pangenome per genomic class, i.e., core, softcore, dispensable, and private.

**Table 2:** *A. thaliana* pangenome graphs characteristics.

| Chromosome | Length | Nodes | Edges | Paths | Steps | A | C | G | N | T |
| --- | --- | --- | --- | --- | --- | --- | --- | --- | --- | --- |
| chr1 | 116770065 | 7906460 | 11365549 | 197 | 540100908 | 36074457 | 20694558 | 20986786 | 3362246 | 35652018 |
| chr2 | 81668410 | 5181104 | 7408132 | 255 | 395411168 | 24615920 | 15403786 | 14887044 | 1713074 | 25048586 |
| chr3 | 93030971 | 6256236 | 8966359 | 222 | 461772242 | 27977964 | 16813683 | 16998921 | 3503434 | 27736969 |
| chr4 | 91053009 | 4992770 | 7189634 | 242 | 467101744 | 26544923 | 17679003 | 16780535 | 2432835 | 27615713 |
| chr5 | 106253880 | 6054673 | 8614824 | 214 | 533961296 | 31628209 | 20057808 | 19495677 | 2336919 | 32735267 |
| Total | 488776335 | 30391243 | 43544498 | 1130 | 2398347358 | 146841473 | 90648838 | 89148963 | 13348508 | 148788553 |

Figure 3 shows the distribution of node lengths across the different genomic classes. Core nodes primarily comprised sequences of length between 2 and 100 bp, while the majority of the nodes included in the softcore, dispensable, and private classes were 1 bp long. Longer nodes, i.e., between 501 and 1,000 bp and longer than 1,000 bp, were mainly found in the dispensable and private classes, while their presence was scarce among the core and softcore classes.

**Figure 3:**
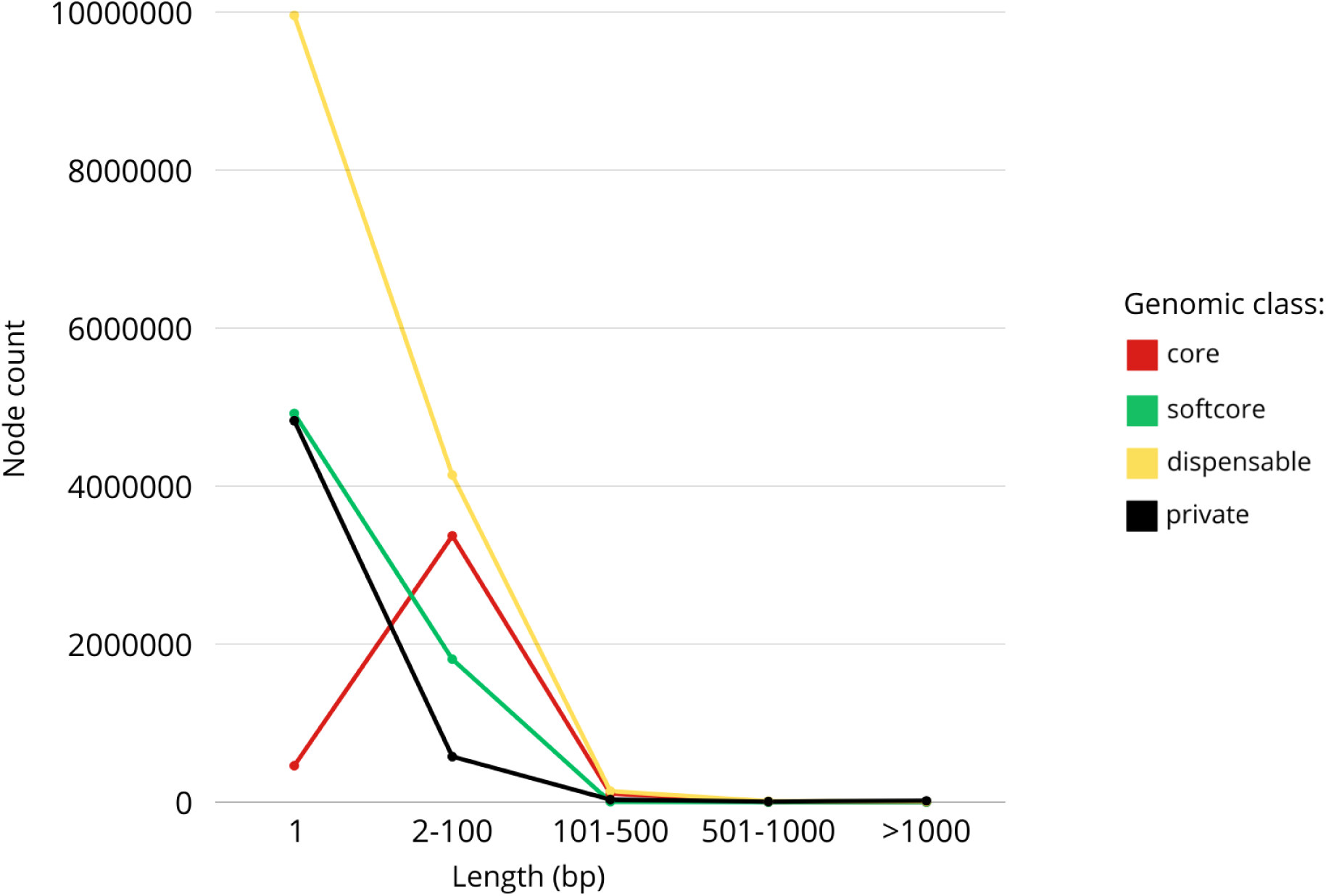
Node length distribution per genomic class in the A. thaliana pangenome.

Figure 4 shows the proportion of nodes divided into genomic classes, i.e., core, softcore, dispensable, and private for each accession. As nodes may be present more than once in a single accession, here only their presence or absence in each accession was considered. As expected, the number of core nodes was equal for all the accessions (3,939,502), while all the other genomic classes showed a certain level of variability in terms of number of nodes per accession. The number of softcore nodes per accession ranged from 6,496,738 to 5,818,444; the number of dispensable nodes per accession ranged from 4,119,053 to 2,942,578; and the number of private nodes per accession ranged from 321,878 to 17. Overall, we found the dispensable class to have the largest range of variability.

**Figure 4:**
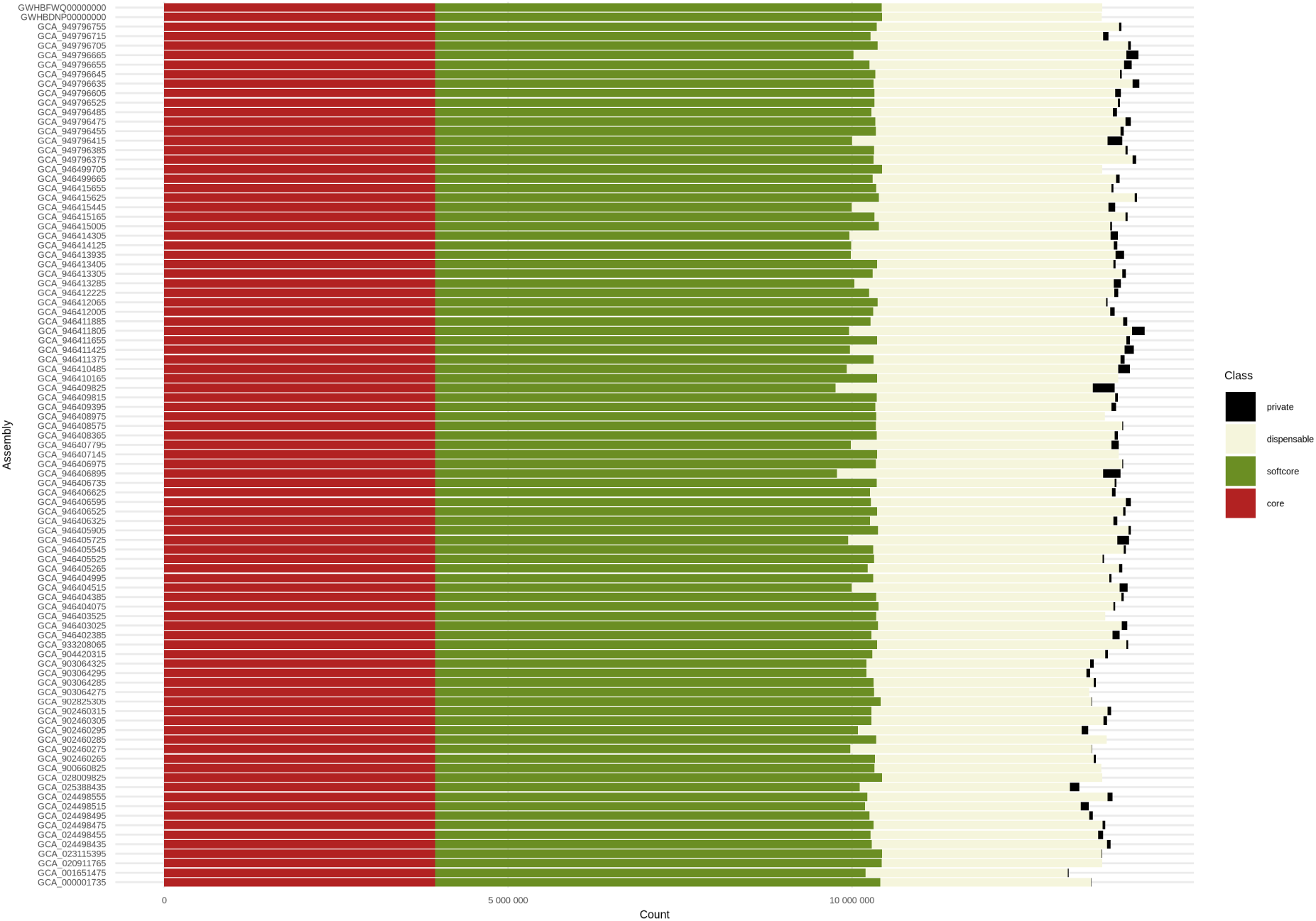
Presence/absence variation of nodes per assembly categorised into genomic classes (i.e., core, softcore, dispensable, and private) in the *A. thaliana* pangenome

Figure 5 shows the similarity tree obtained from the Jaccard distance matrix calculated between the accessions’ sequences. To highlight potential geographic-specific clusters, we coloured the tree tips according to the accessions’ country of origin. The cophenetic correlation coefficient, which was equal to 0.8, confirmed the tree’s robustness. The plot shows some country-specific clusters, but the Mantel test indicated a correlation coefficient of 0.05996592 and a simulated p-value of 0.1859, which means that there was only a weak and not statistically significant link between the genetic distance based on sequence divergence and geographical distance.

**Figure 5:**
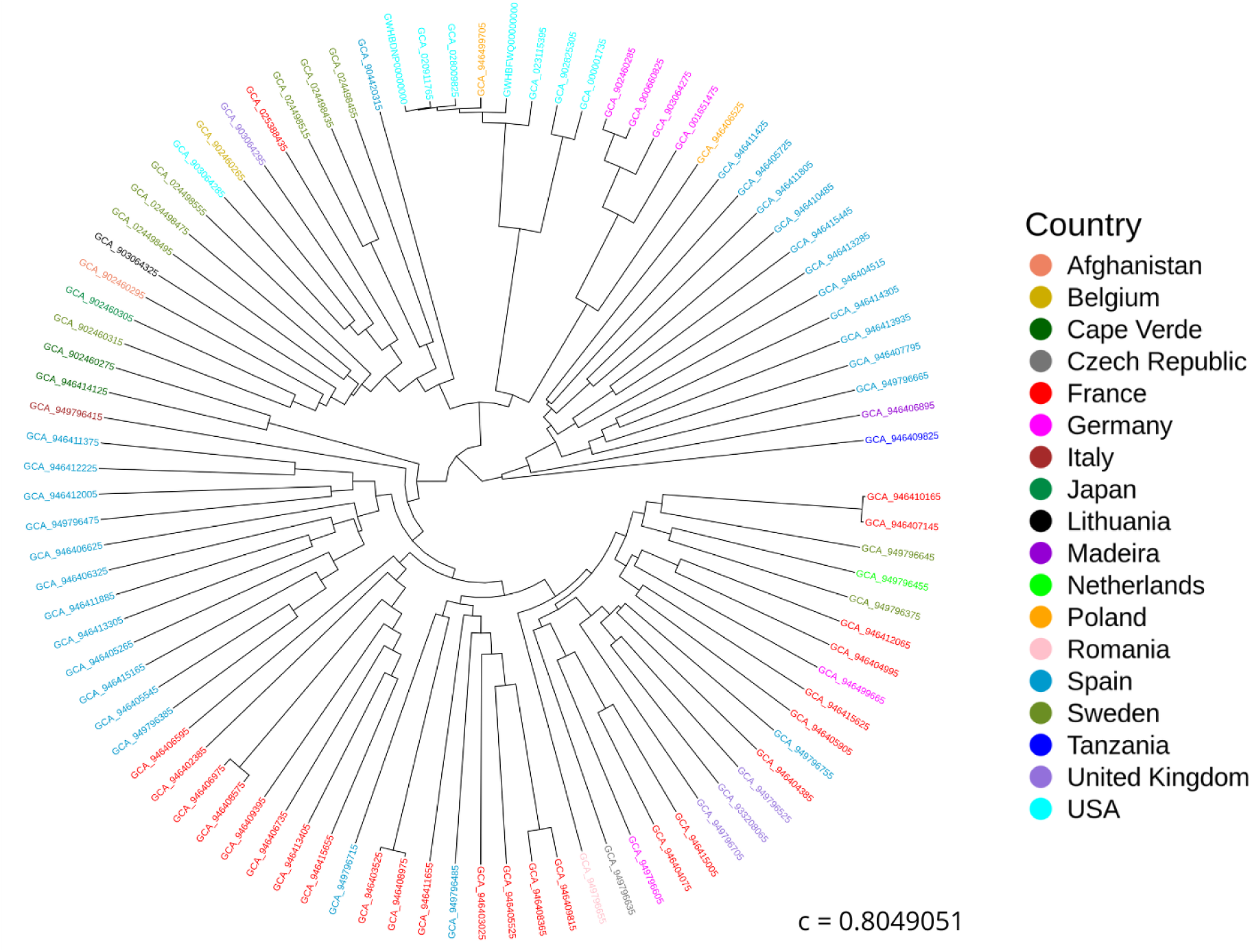
Similarity tree based on the Jaccard distance between assemblies calculated on the sequences (i.e., nodes). “c” represents the cophenetic correlation coefficient.

### 3.3 Gene-based pangenome

#### 3.3.1 Reference-based pangenome annotation

In the first step of the analysis, the genes annotated on the reference genome TAIR10.1 were injected into the pangenome graph and mapped across the 93 assemblies. We were able to map to the pangenome the 86.3% of the reference genes (i.e., 28,512) with 95% of estimated identity and coverage. A total of 9,487 genes were classified as core; 11,518 were classified as softcore; 7441 were classified as dispensable; and 66 were classified as private. The list of unmapped genes is shown in Supplementary Materials Table S3.

Of these genes, we recorded the copy number variation. The overall percentage of genes showing copy number variation was 2.8% (791), of which 112 were core genes, 334 softcore genes, and 345 dispensable genes, while no private gene showed copy number variation. The complete list of genes showing copy number variation in at least one accession is shown in Supplementary Materials Table S4.

Figure 6 shows the distribution of mapped reference genes considering copy number variation for each assembly, separated into genomic classes, i.e., core, softcore, dispensable, and private. The core and private genomic classes exhibited a narrow range of variability. The number of core genes varied from 9,524 to 9,500. As expected, the reference genome TAIR10.1, specifically GCA 000001735, contained the highest number of private genes (25). All injected reference genes were expected to map to the reference genome and, eventually, also to other genomes, meaning that none of these genes were expected to be classified as private for non-reference genomes. However, 41 reference genes were mapped on 28 non-reference assemblies with coverage and estimated identity equal to at least 95%, and not on TAIR10.1 (Supplementary Materials Table S5). 64 assemblies did not show private reference genes. All reference private genes were present in a single copy. The maximum number of private genes mapped on a non-reference assembly was 4, observed in GCA 946406895 and GCA 949796415. The number of softcore genes per assembly ranged from 11,717 to 9,336, and the number of dispensable genes per assemblies varied from 7,491 to 2,301. Overall, we found the dispensable class to have the largest range of variability.

**Figure 6:**
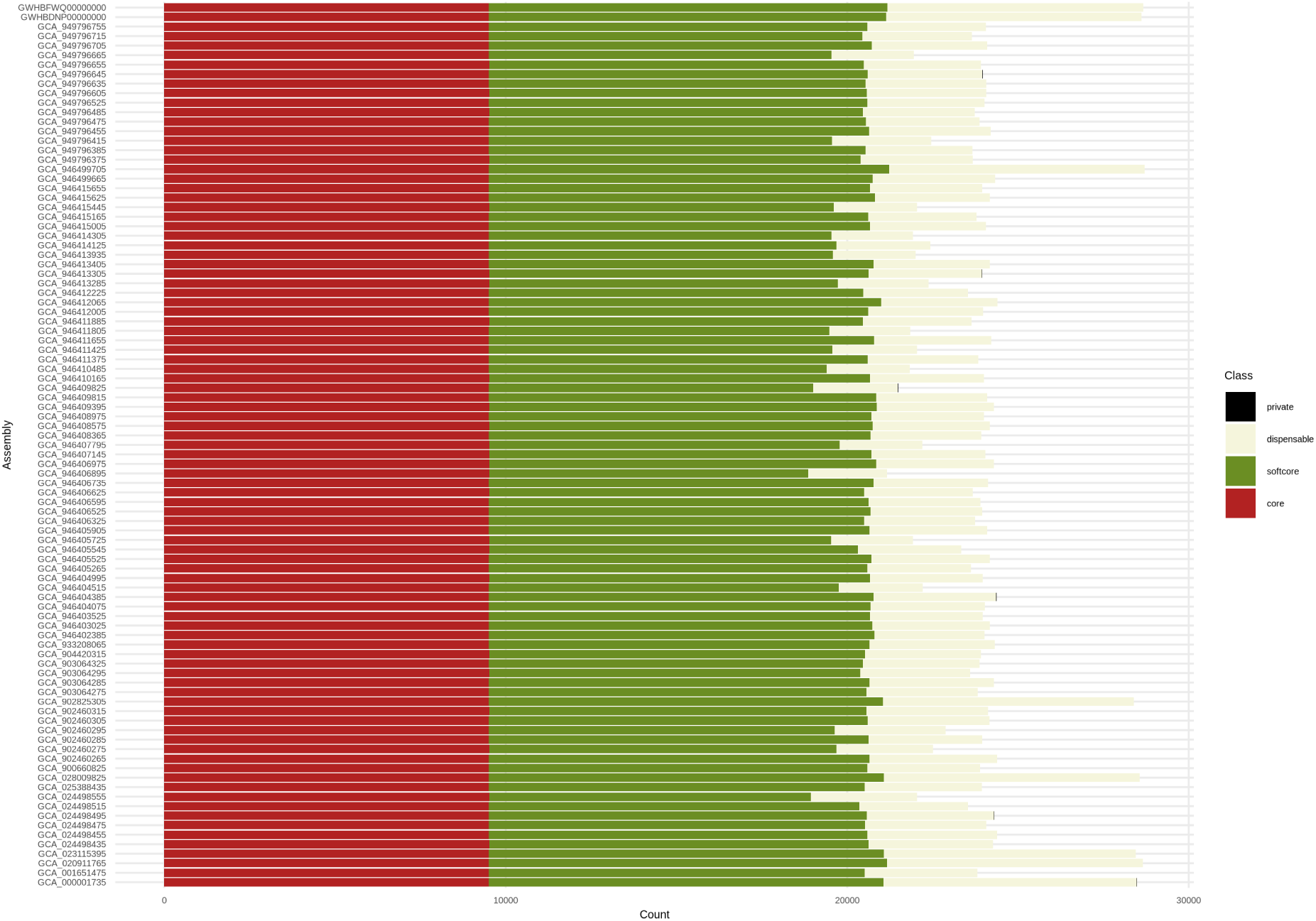
Copy number variation per assembly of the reference genes mapped across the 93 assemblies with 95% of estimated identity and coverage.

#### 3.3.2 Final pangenome annotation

In the second step of the analysis, we extracted from the 92 non-reference assemblies the sequences that were not shared with TAIR10.1. The non-reference sequences were annotated with the approach Diamond + Exonerate as described in Materials and Methods and processed together with the results obtained from the reference derived annotations. With this approach, we were able to map additional 2,505 loci, for a final number of annotated loci equal to 31,017. We annotated 23,255 to 28,397 loci per assembly.

In this phase of the analysis, we only considered presence or absence of loci (Presence Absence Variation, PAV), and CNVs were recorded as a binary variation.

Figure 7 illustrates the relative gene composition of the *A. thaliana* pangenome. We categorised 36% of genes as core (11,152), 34.7% as softcore (10,754), 28.1% as dispensable (8,731), and 1.2% as private (380).

**Figure 7:**
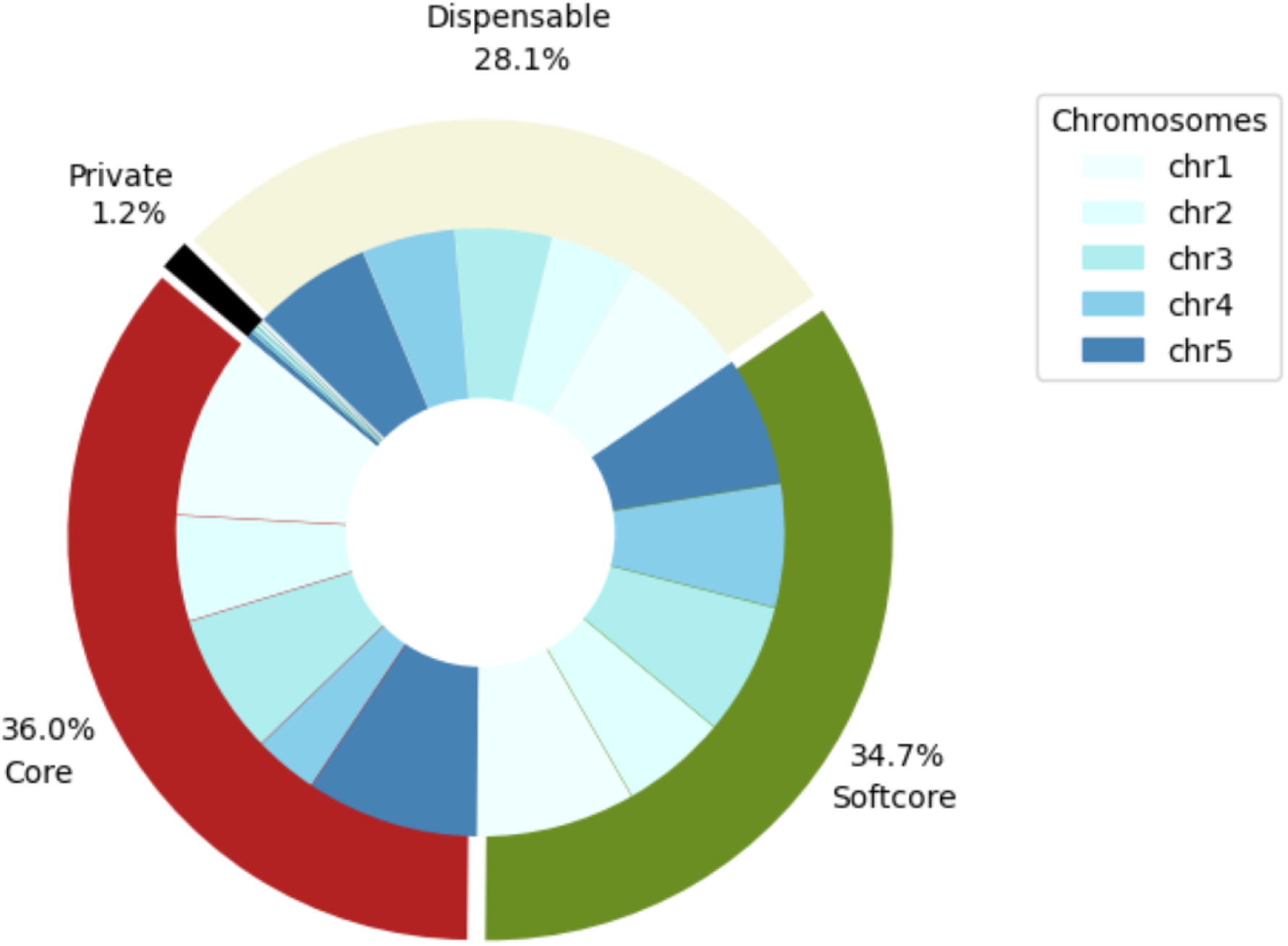
Relative gene content of the A. thaliana pangenome per genomic class, i.e., core, softcore, dispensable, and private.

Figure 8 shows the proportion of genes divided into genomic classes for each assembly. As expected, the number of core genes was equal for all the accessions (11,152), while all the other genomic classes showed a certain level of variability in terms of number of genes per accession. The number of softcore genes per assembly varied from 10,676 to 8,895; the number of dispensable genes per assembly ranged from 6,557 to 2,921; and the number of private genes per assembly varied from 45 to 0. Overall, we found the dispensable class to have the largest range of variability.

**Figure 8:**
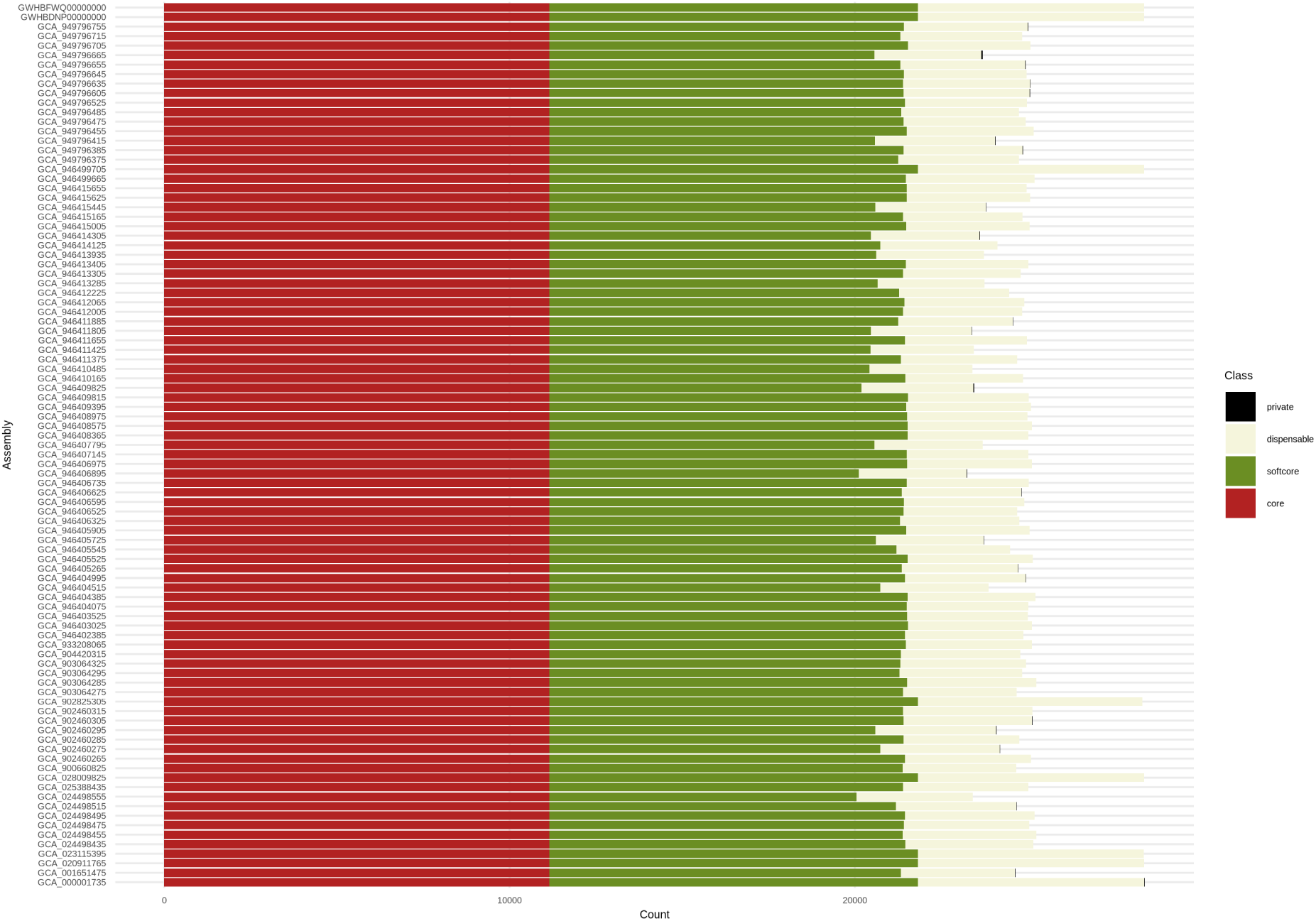
Presence/absence variation of genes per assembly categorised into genomic classes (i.e., core, softcore, dispensable, and private) in the *A. thaliana* pangenome.

We defined as new genes those annotated on the non-reference sequences that did not have a corresponding annotation on TAIR10.1 in the official conversion table TAIR to UniProt. They are shown in Supplementary Materials Table S6.

Of the 250 new genes, 9 were classified as softcore genes (IDs: UniRef100 P0CAY3, UniRef100 Q9LMS5, UniRef100 P0CAY1, UniRef100 Q9LJ82, UniRef100 Q9M8J5, UniRef100 Q84W54, UniRef100 Q93ZE5, UniRef100 F2Q9V4, and UniRef100 P0DN92), 187 as dispensable, and 42 as private.

234 genes identified as new (93,6%) were attributed to *A. thaliana*, while 16 (6.4%), reported in Table 3, were attributed to other species.

**Table 3:**
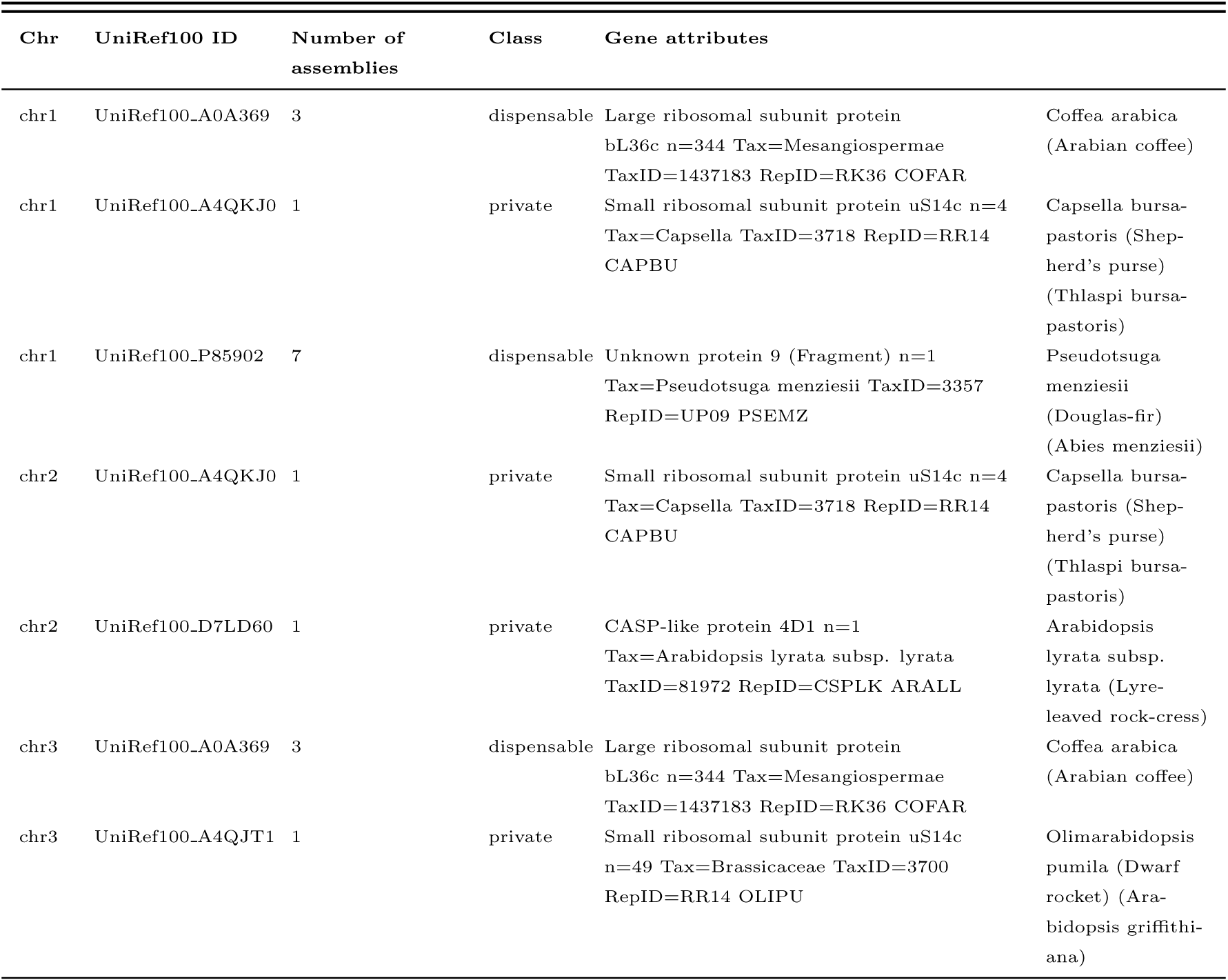
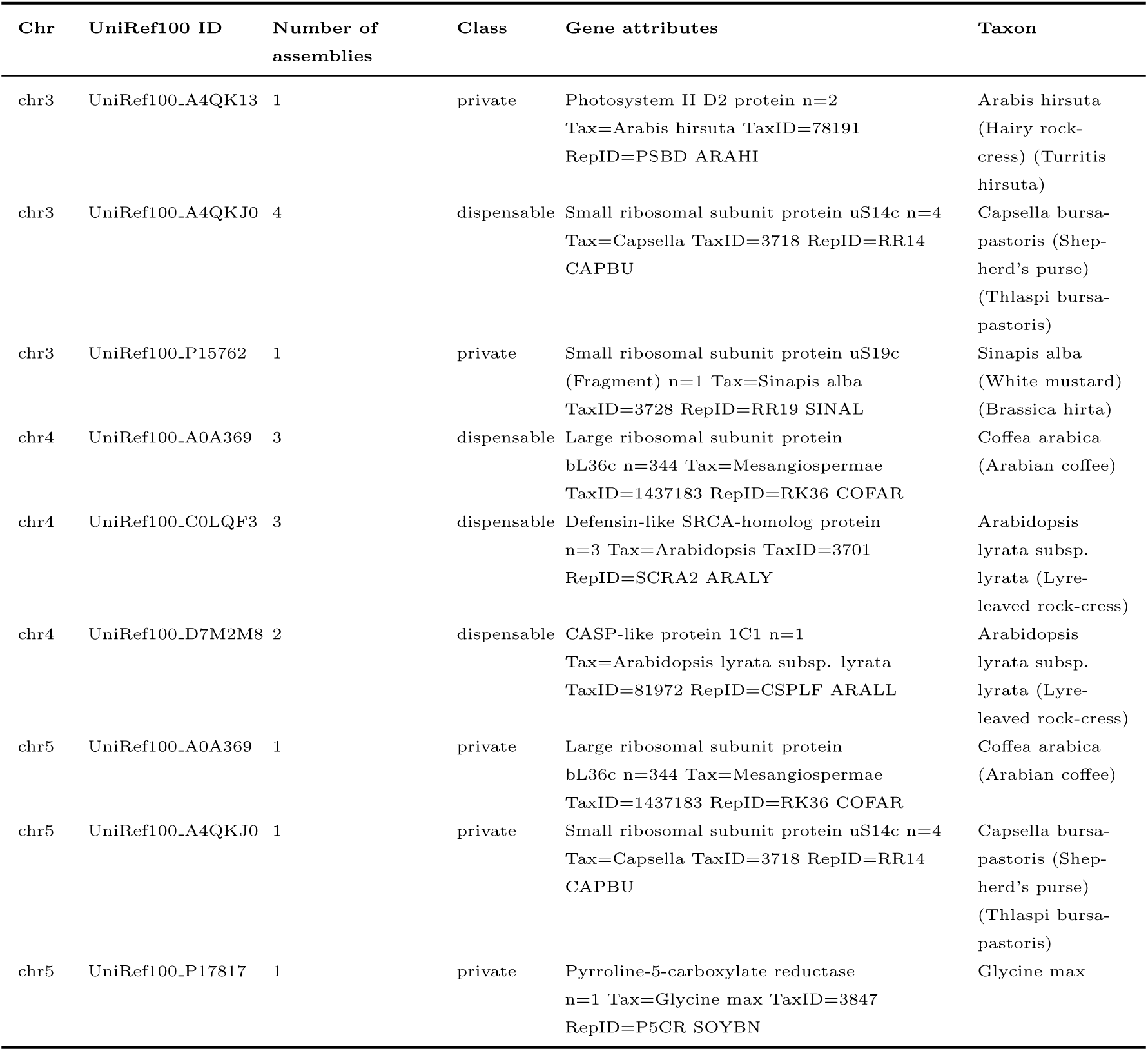
New genes found in non-reference sequences attributed to species different from *A. thaliana*.

We classified 9 of the 16 new genes attributed to other species as private, while we classified the remaining 7 as dispensable and found them in 2 to 7 accessions.

We found UniRef100 A0A369 locus attributed to *Coffea arabica* in a total of 10 accessions. This locus mapped in non collinear positions (on chromosome 1 for GCA 946406625, GCA 946406735, and GCA 946413305; on chromosome 3 for GCA 025388435, GCA 902460315, and GCA 933208065; on chromosome 4 for GCA 946403025, GCA 946405905, and GCA 946412065; and on chromosome 5 for GCA 946409825), and therefore considered as a different gene when found on different chromosomes.

The locus attributed to *Capsella bursa-pastoris* was found in 7 accessions in total, but on different chromosomes (chromosome 1 for GCA 946411885; chromosome 2 for GCA 001651475; chromosome 3 for GCA 025388435, GCA 902460305, GCA 933208065, and GCA 946409825; and chromosome 5 for GCA 946409825), and therefore considered as a different gene when found on different chromosomes.

The gene translocation events between chromosomes are described in Supplementary Materials Table S7. In this table we also included genes that are annotated on the organelles in TAIR10.1, i.e., mitochondrion and chloroplast.

We observed a total of 344 translocations, of which 212 were classified as dispensable and 132 as private. No core or softcore translocations were detected.

Figure 9 illustrates the distribution of translocations between chromosomes. Chromosome 1 received translocations primarily from chromosomes 2 and 5. Chromosome 2 showed translocations mainly coming from chromosome 3, and an important presence of genes from the mitochondrion. Chromosome 3 exhibited the highest number of total translocations, the majority of which involved genes coming from chromosome 2 and the mitochondrion. In chromosome 4, a balanced distribution of translocations from all the chromosomes was detected, along with genes coming from the chloroplast. Chromosome 5 showed the highest number of nuclear translocations, mainly involving chromosome 1 and 2, and genes coming from both organelles.

**Figure 9:**
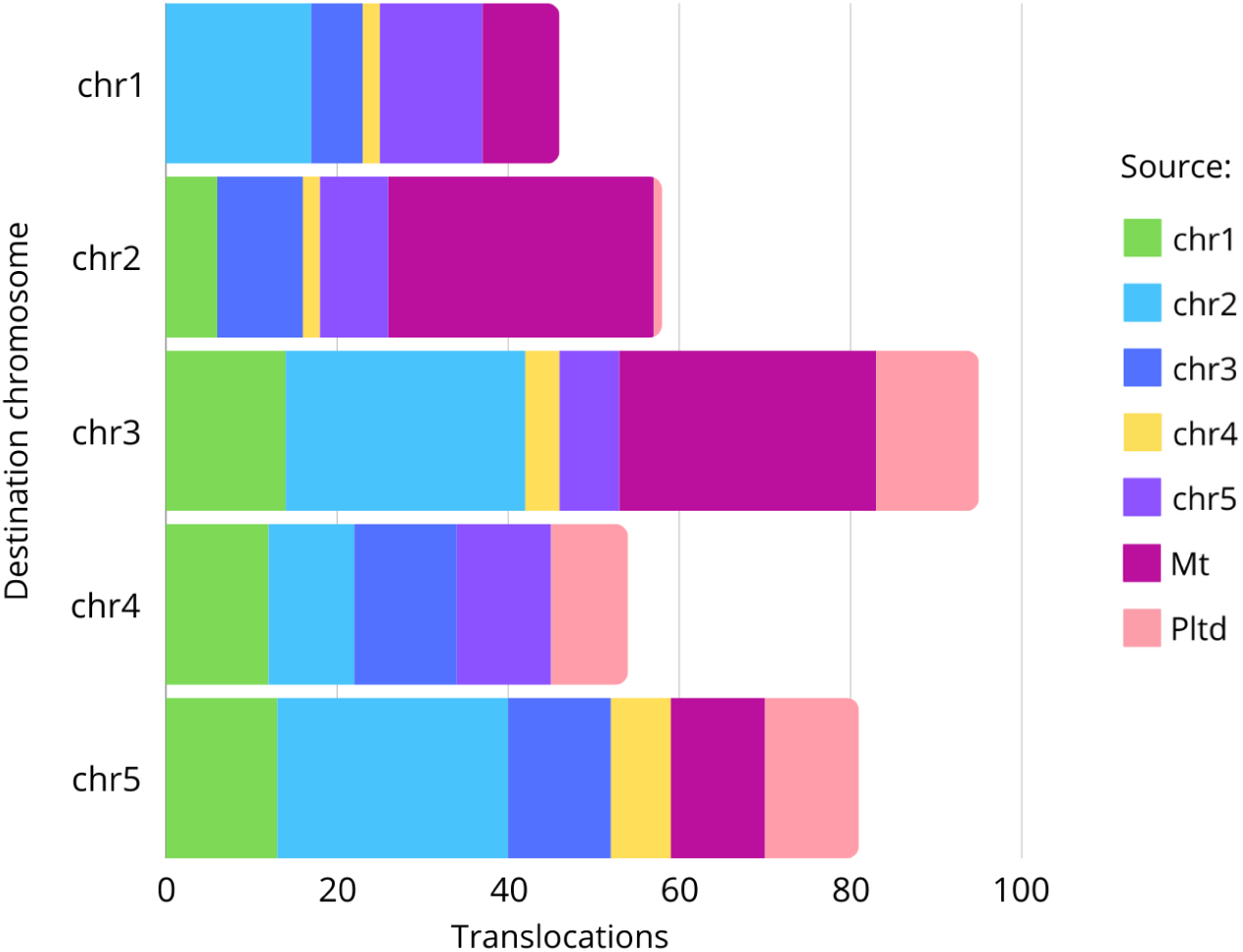
Distribution of nuclear gene translocations and organellar gene insertions per chromosome in the A. thaliana pangenome.

We explored the gene ontology enrichment of the genes classified into core and variable (i.e., softcore, dispensable, and private) using TopGO.

Core genes were enriched for gene ontologies related to primary and essential biological processes, including, for example, developmental processes, multicellular organismal processes, reproduction, and biological regulation. Variable genes (the ensemble of softcore, dispensable and private) were significantly enriched for genes involved in catabolic processes of several cellular compounds, including, for example, aromatic and organitrogen compounds and RNA (Supplementary Materials Figure S2 and Figure S3).

The similarity tree in Figure 10 was obtained from the Jaccard distance matrix calculated on gene PAVs among the accessions. The tips of the tree were coloured according to the country of origin of the accessions to highlight possible geographic-specific clusters. The cophenetic correlation coefficient (c = 0.8) validated the tree’s robustness. The Mantel test found a correlation coefficient of 0.2017449 and a simulated p-value of 0.0073. This result indicated a moderately positive and statistically significant correlation between the genetic distance in terms of gene PAVs and the geographic distance between the accessions.

**Figure 10:**
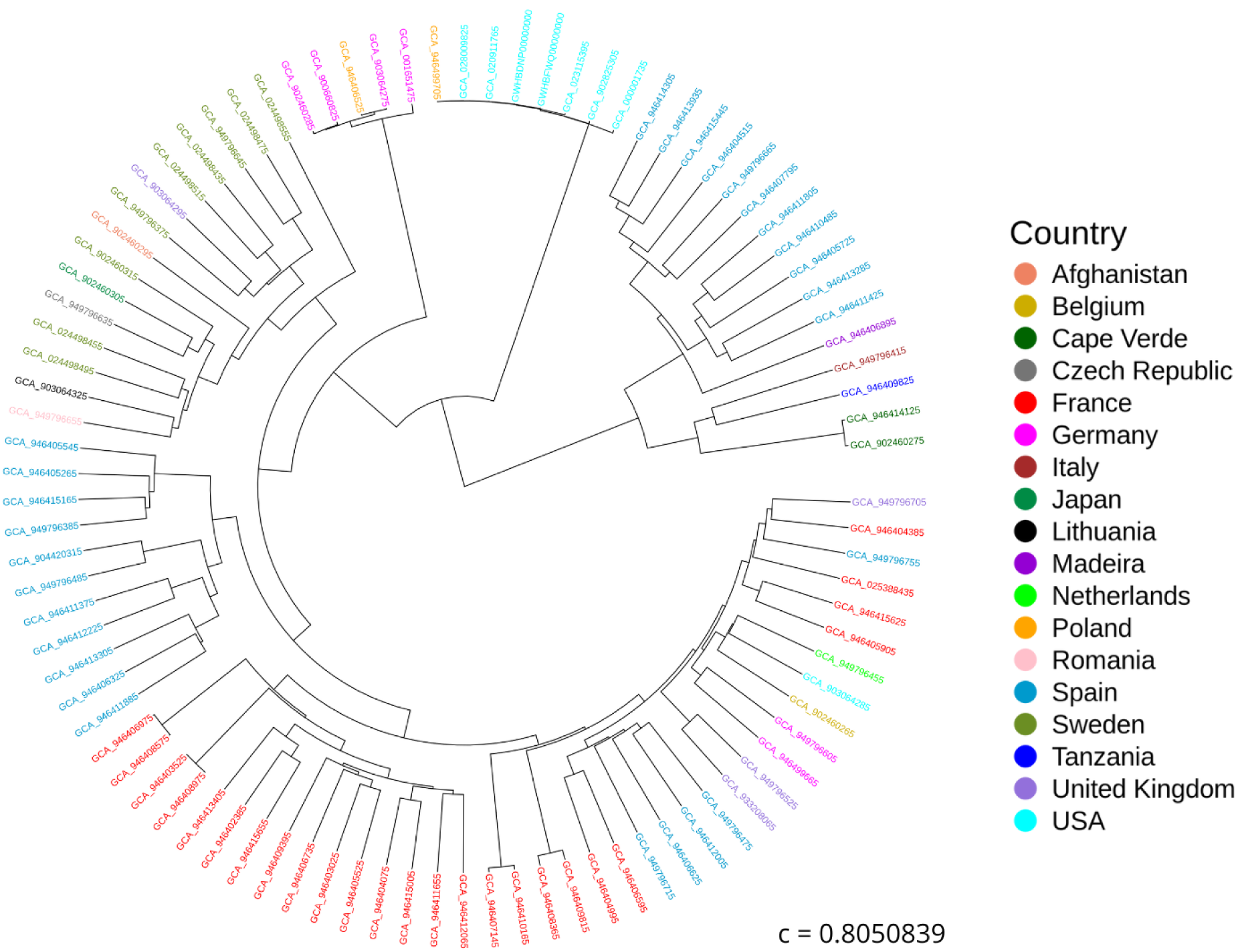
Similarity tree based on the Jaccard distance between assemblies calculated on the gene PAVs. “c” represents the cophenetic correlation coefficient.

To estimate the pangenome growth in terms of gene content, we calculated gene counts when different size of pangenome is considered, as detailed in Materials and Methods. As expected, the total number of genes increases with the number of assemblies considered. However, although the slope of the curve decreases, it seems that a plateau was not reached. The number of core genes showed a trend opposite to that of softcore and dispensable genes. The private genes decreased with the number of assemblies in the pangenome (Figure 11).

**Figure 11:**
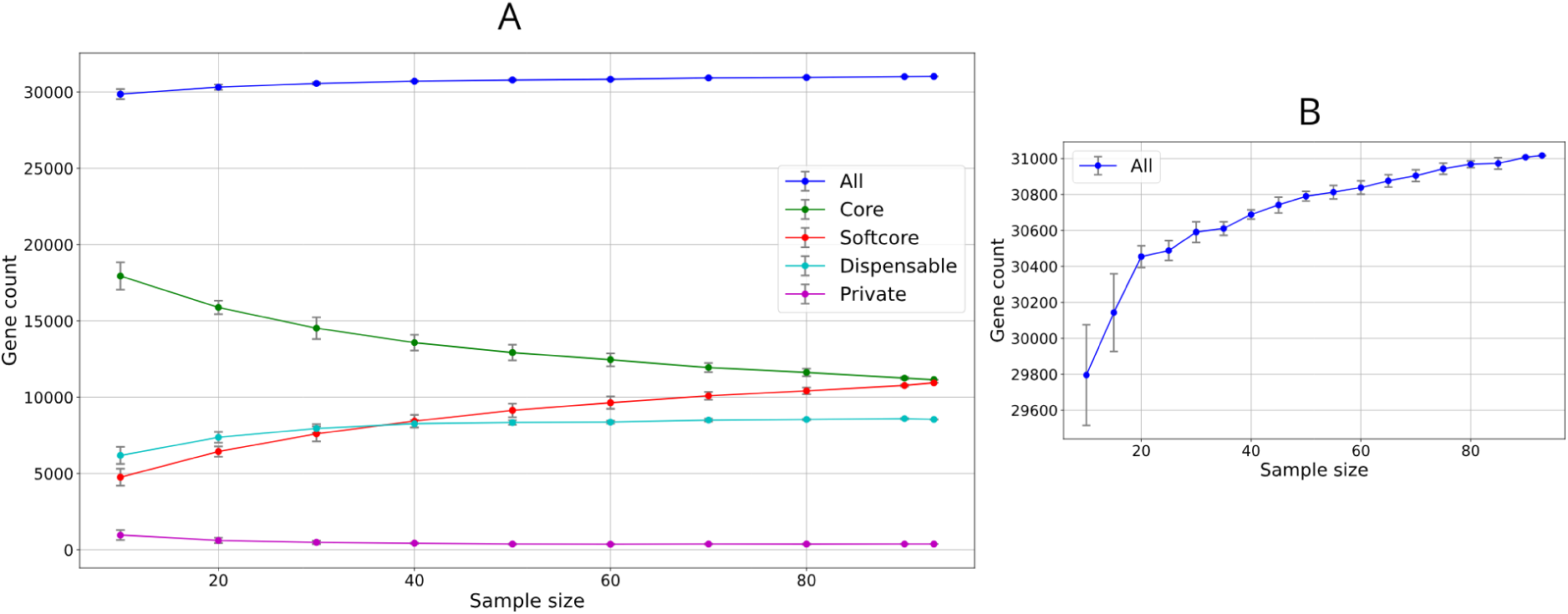
Simulated A. thaliana pangenome pangenome growth curve in terms of gene content. A shows all the genomic classes, while B shows only the cumulative curve including all classes together.

### 3.4 Pseudogene-based pangenome

#### 3.4.1 Reference-based pangenome annotation

Initially, the pseudogenes annotated on the reference genome TAIR10.1 by Mascagni et al. (2021) [35] were injected into the pangenome graph and mapped across the 93 assemblies. We successfully mapped 96% of the reference pseudogenes (i.e., 3,555) to the pangenome graph with 95% estimated identity and coverage. Of these, 186 pseudogenes were designated as core; 640 as softcore; 699 as dispensable; and 2,030 as private.

We reported the copy number variation of these pseudogenes. The total percentage of pseudogenes exhibiting copy number variation was 19% (668), comprising 49 core pseudogenes, 204 softcore pseudogenes, 190 dispensable pseudogenes, and 225 private pseudogenes. The whole list of pseudogenes exhibiting copy number variation in at least one accession are shown in Supplementary Materials Table S8.

Figure 12 illustrates the distribution of mapped reference pseudogenes, incorporating copy number variation throughout each assembly, categorised into genomic classes, i.e., core, softcore, dispensable, and private.

**Figure 12:**
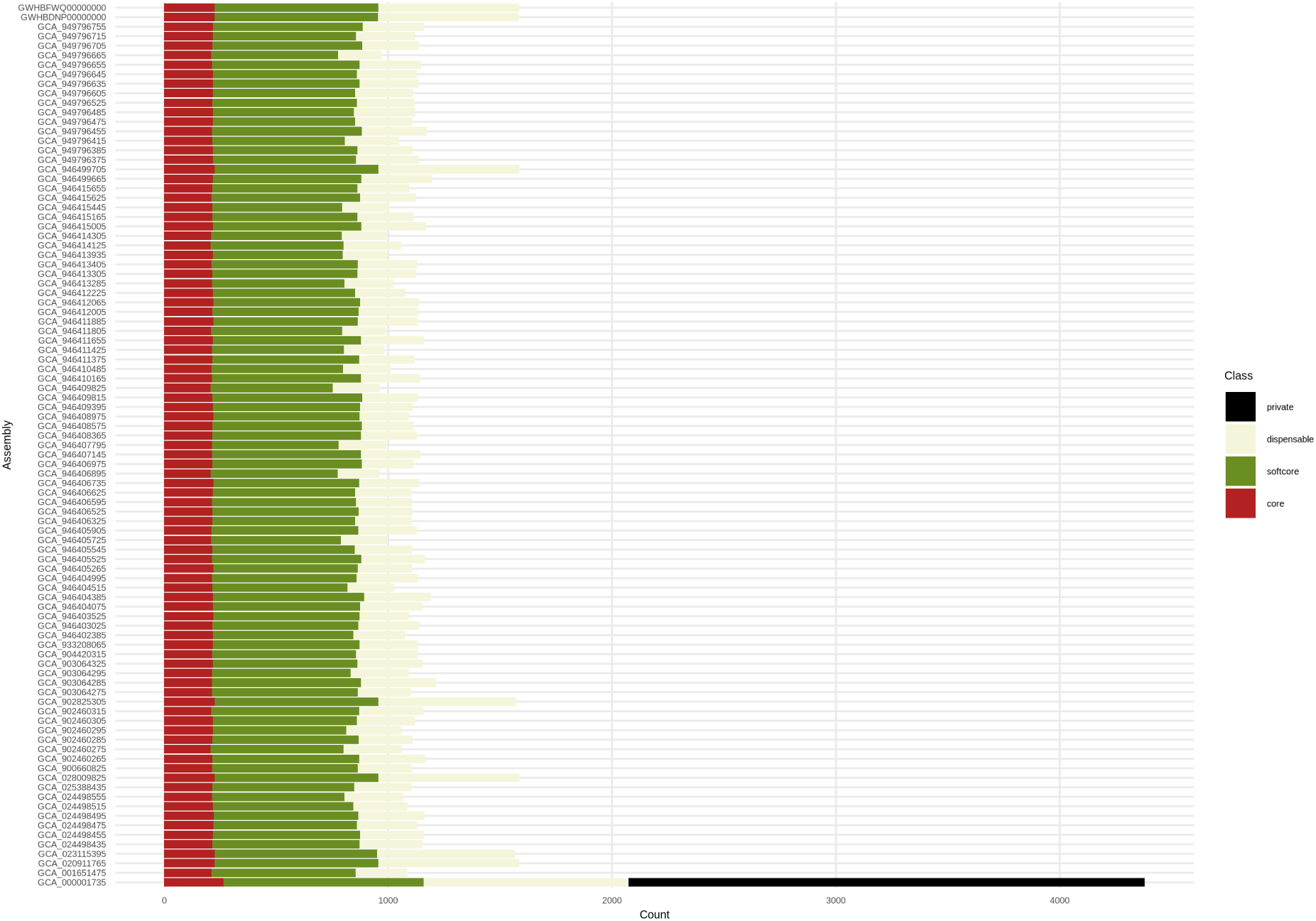
Copy number variation per assembly of the reference pseudogenes mapped across the 93 assemblies with 95% of estimated identity and coverage.

The accession GCA 000001735 (i.e., TAIR10.1) carried the highest quantity of core (264), softcore (895), dispensable (915), and private (2,306) pseudogenes. This result was expected, but the number of copy number variation on the other 92 assemblies might be biased by the filter applied. In fact, the pseudogenes that mapped on to the non-reference accessions were removed when they were lacking stop codons or exhibited coverage equal to 100%, as we could not affirm with certainty that the feature was not complete and functional on a given assembly. This might have led to an underestimation of the quantity of pseudogenes found in the non-reference assemblies.

The accession GCA 946409825 displayed the lowest quantity of core (207) and softcore pseudogenes (545). The accession GCA 946411425 carried the lowest number of dispensable pseudogenes (183). 91 accessions possessed no private pseudogenes. We expected to find no private pseudogenes in any of the non-reference assemblies, but we found that one pseudogene, originating from the parent gene AT3G17320, was mapped to the chromosome 4 of the assembly GCA 946407795 and not to the reference TAIR10.1 and it was present with a single copy. Overall, we found the private class to have the largest range of variability, although, as explained, this interpretation suffers from a bias.

#### 3.4.2 Final pangenome annotation

In the second phase of the study, we retrieved sequences from the 92 non-reference assemblies that were not present in TAIR10.1. The non-reference sequences were annotated using the Diamond + Exonerate method and processed alongside the results from the reference-derived annotations to generate the final statistics. This approach enabled us to map an additional 20,339 pseudogenes, resulting in a total of 23,894 identified pseudogenes. We annotated between 1,634 and 6,576 pseudogenes per assembly.

During this stage of the analysis, we just considered the presence or absence of the pseudogenes, ignoring the copy number variations.

Figure 10 depicts the relative pseudogene content of the *A. thaliana* pangenome. We classified the 0.9% of pseudogenes as core (216), 3.9% as softcore (939), 81.9% as dispensable (19,578), and 13.2% as private (3,161).

**Figure 13:**
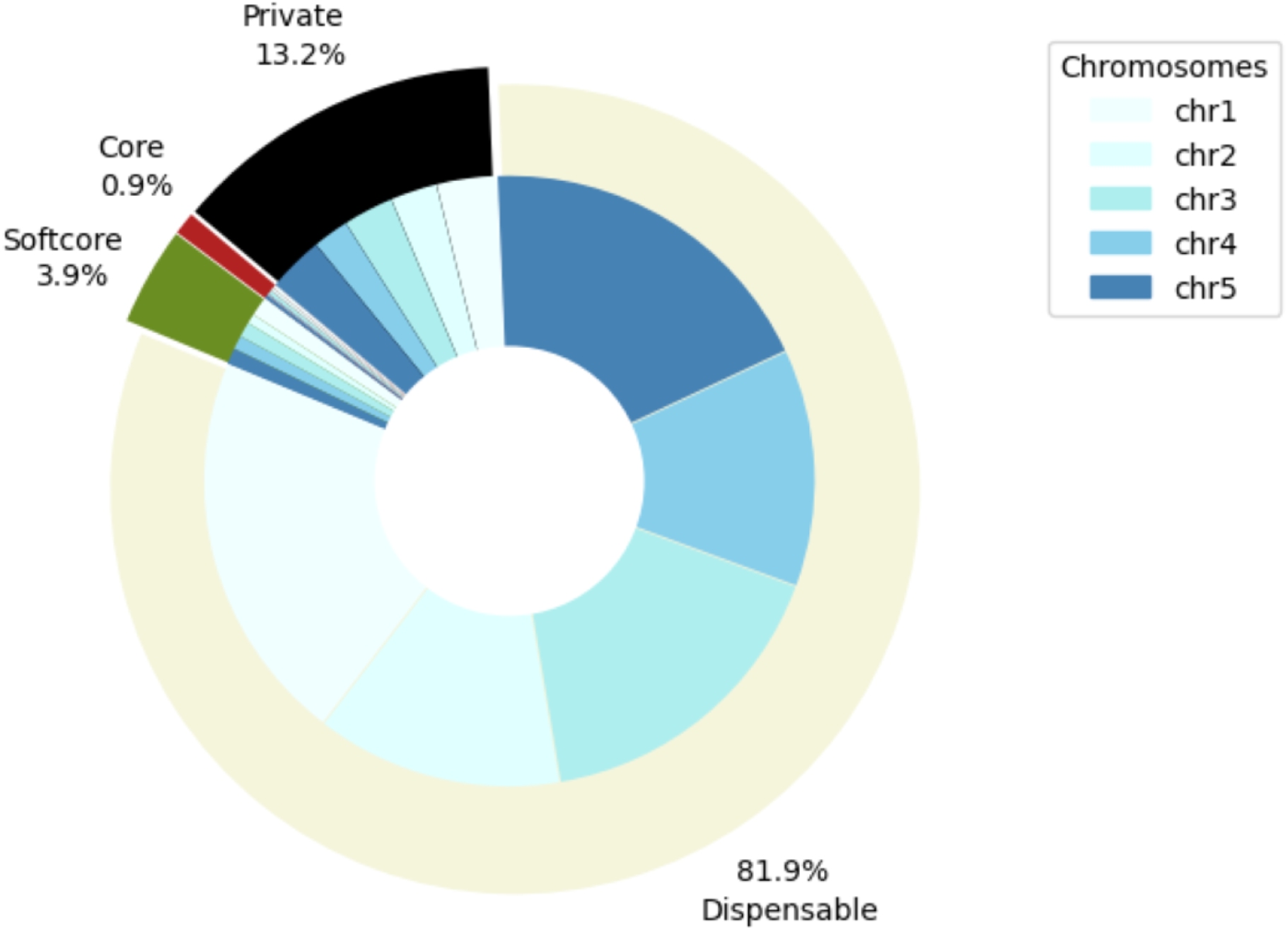
Relative pseudogene content of the A. thaliana pangenome per genomic class, i.e., core, softcore, dispensable, and private.

Figure 14 illustrates the distribution of pseudogenes categorised by genomic classes for each assembly. As expected, the number of core pseudogenes was consistent across all accessions (216), but the other genomic classes exhibited diversity in pseudogenes count per accession. The number of softcore pseudogenes per accession ranged from 908 to 662, and the number of dispensable pseudogenes per assembly varied from 5,444 to 752. The reference genome TAIR10.1 (i.e., GCA 000001735) exhibited the highest number of private pseudogenes. However, this result could still suffer from the bias described above, as the non-reference sequence results of the second phase were added to those obtained from the first stage of the analysis. A total of 10 accessions showed no private pseudogenes. Overall, we observed the greatest variability range for the dispensable class.

**Figure 14:**
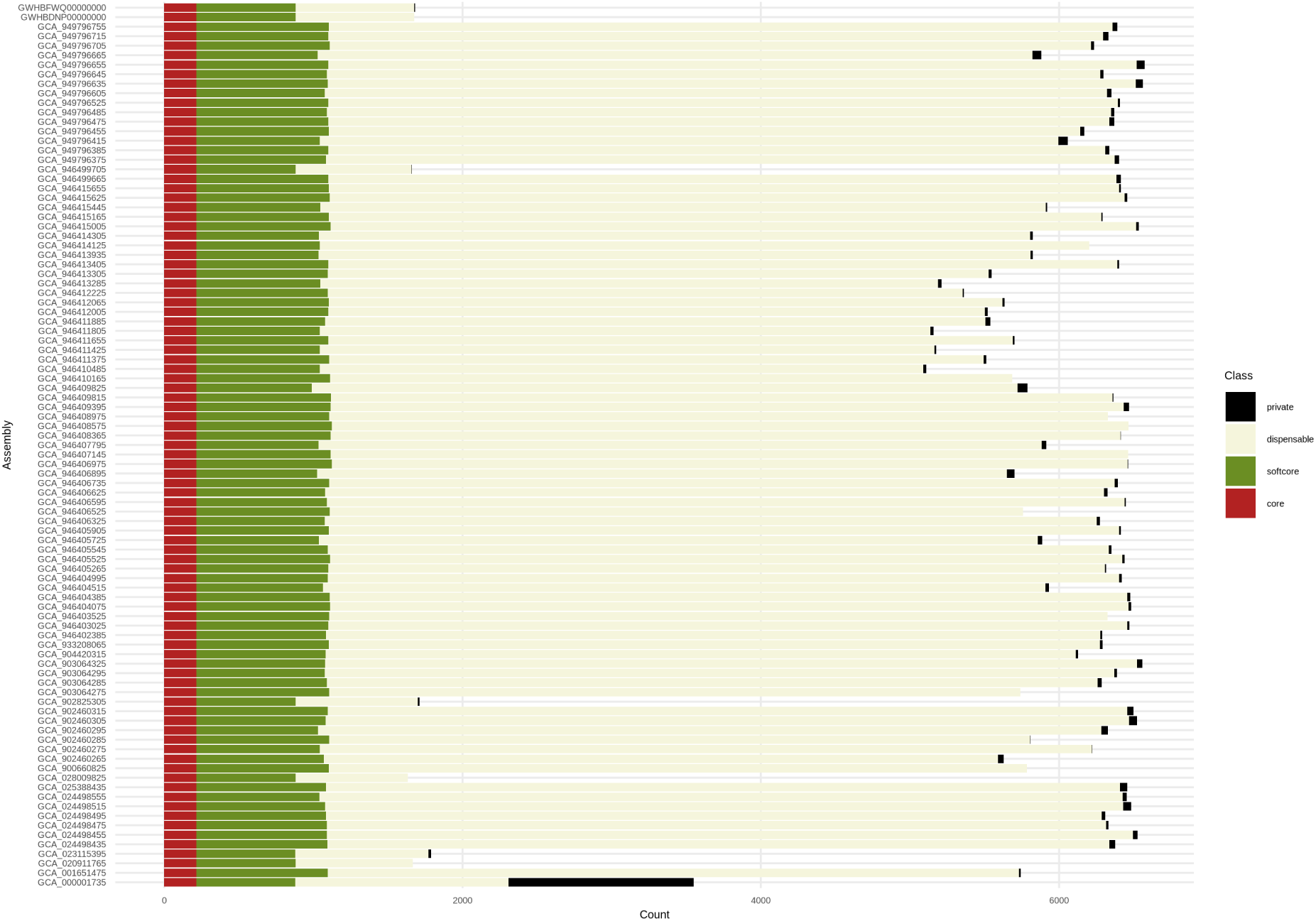
Presence/absence variation of pseudogenes per assembly categorised into genomic classes (i.e., core, softcore, dispensable, and private) in the *A. thaliana* pangenome.

We classified as new pseudogenes those annotated on the non-reference sequences that lacked a corresponding annotation in TAIR10.1 for the corresponding parent gene, according to the official conversion table TAIR to UniProt. The information is shown in Supplementary Materials Table S9.

As expected, no core pseudogenes were found among the new pseudogenes. Of the 460 new pseudogenes, 20 were classified as softcore, 383 as dispensable, and 57 as private.

The 90.1% of the new pseudogenes were attributed to the *A. thaliana* species, while the 9.3%, reported in Table 4, were attributed to other species.

**Table 4:**
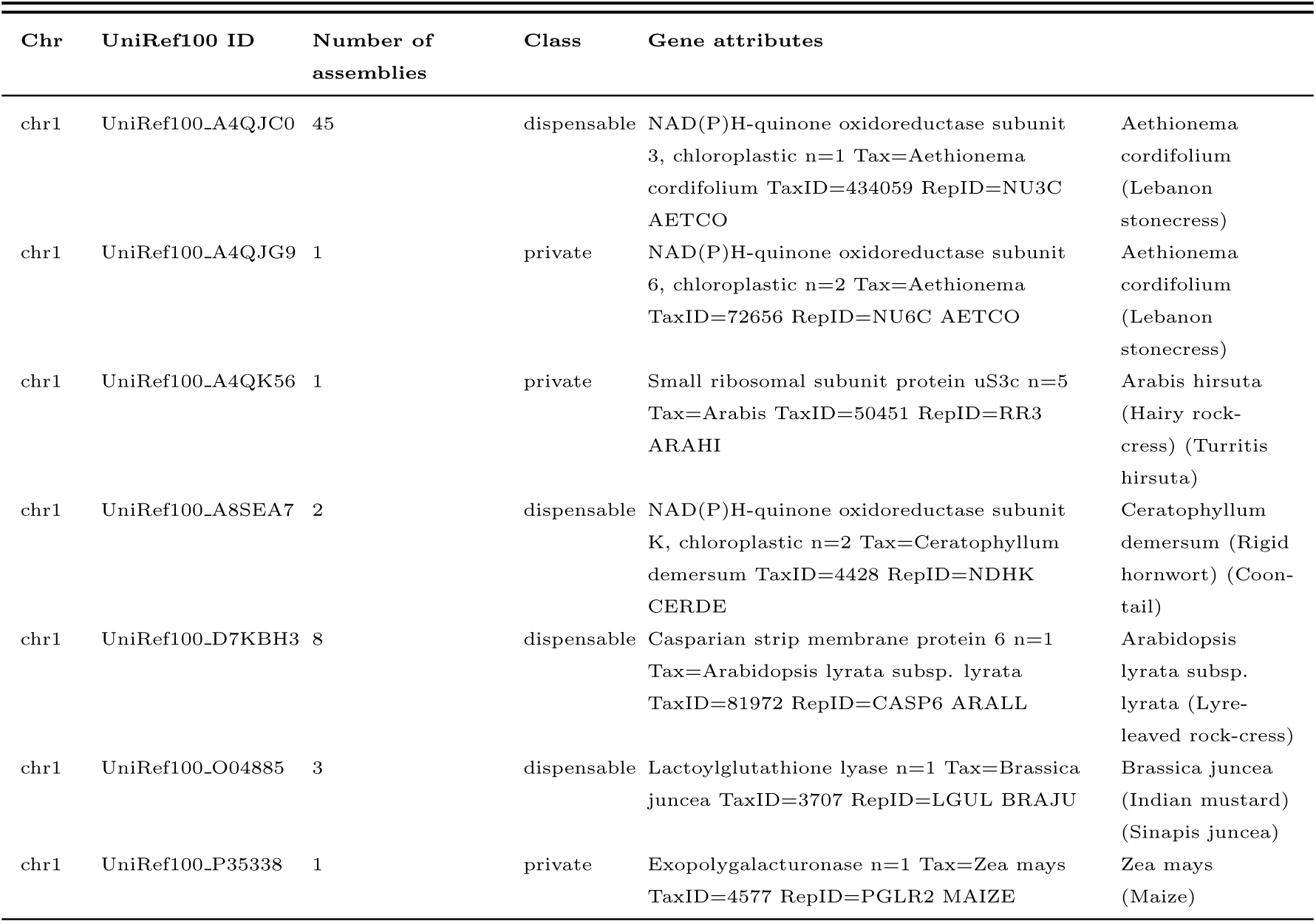
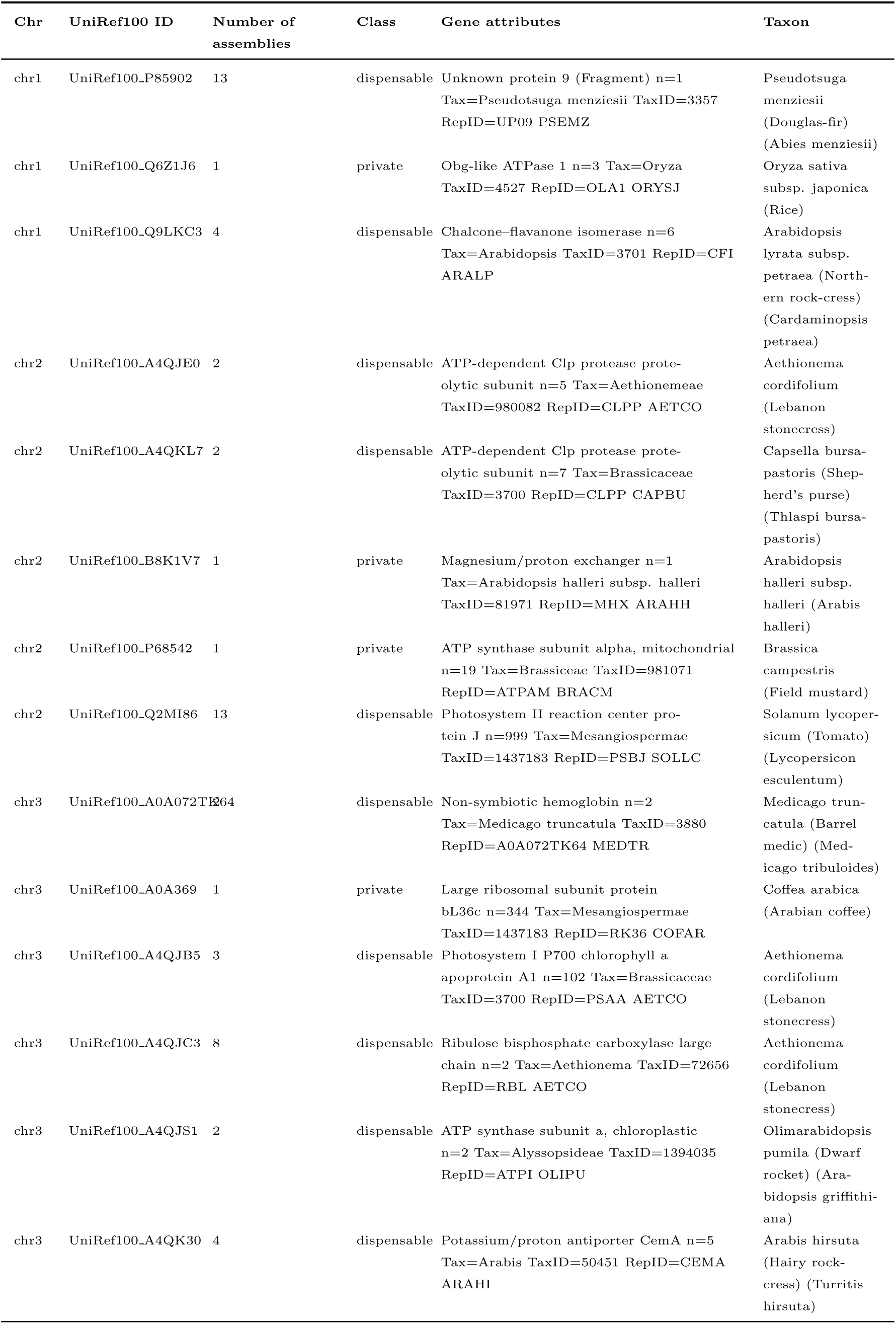
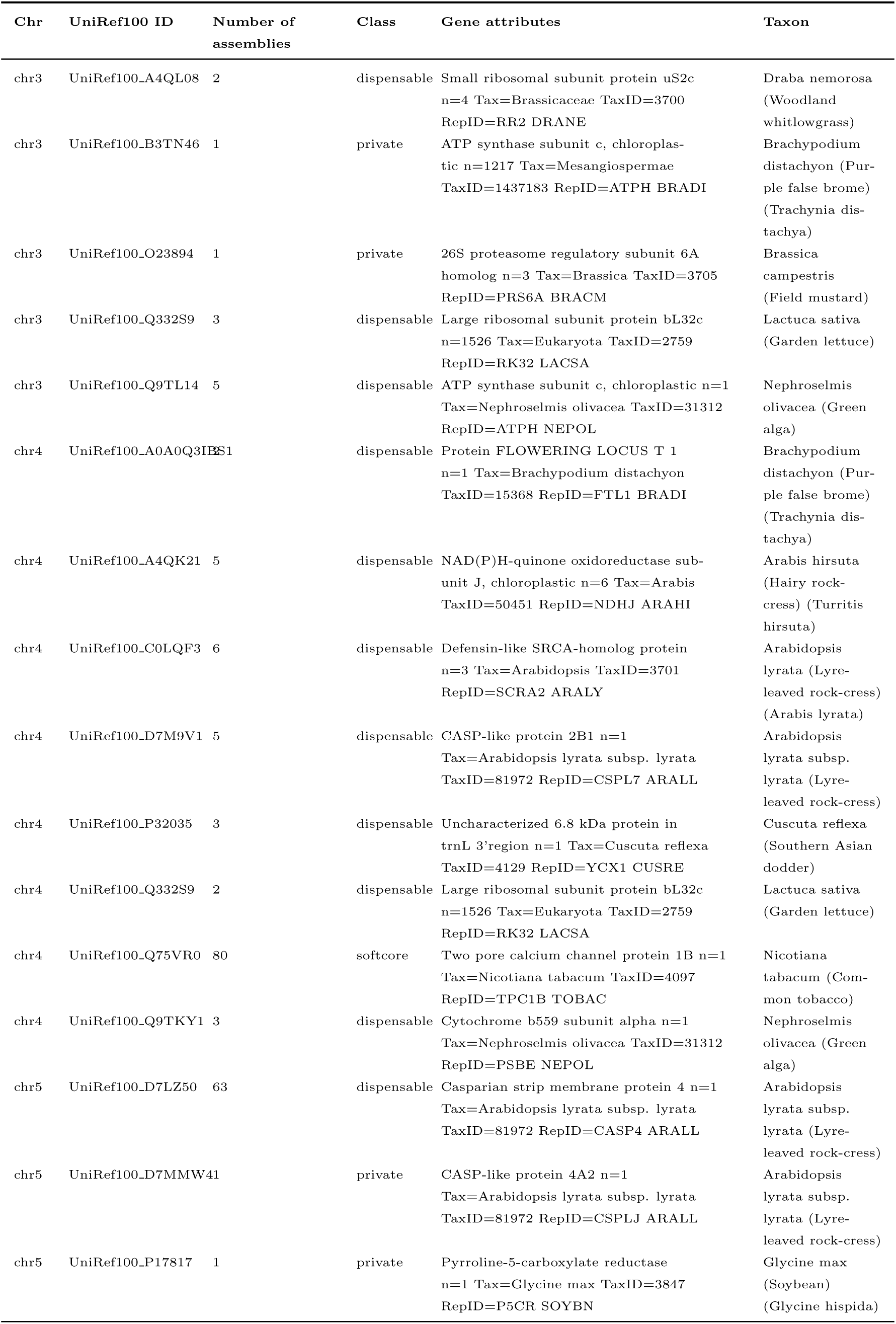
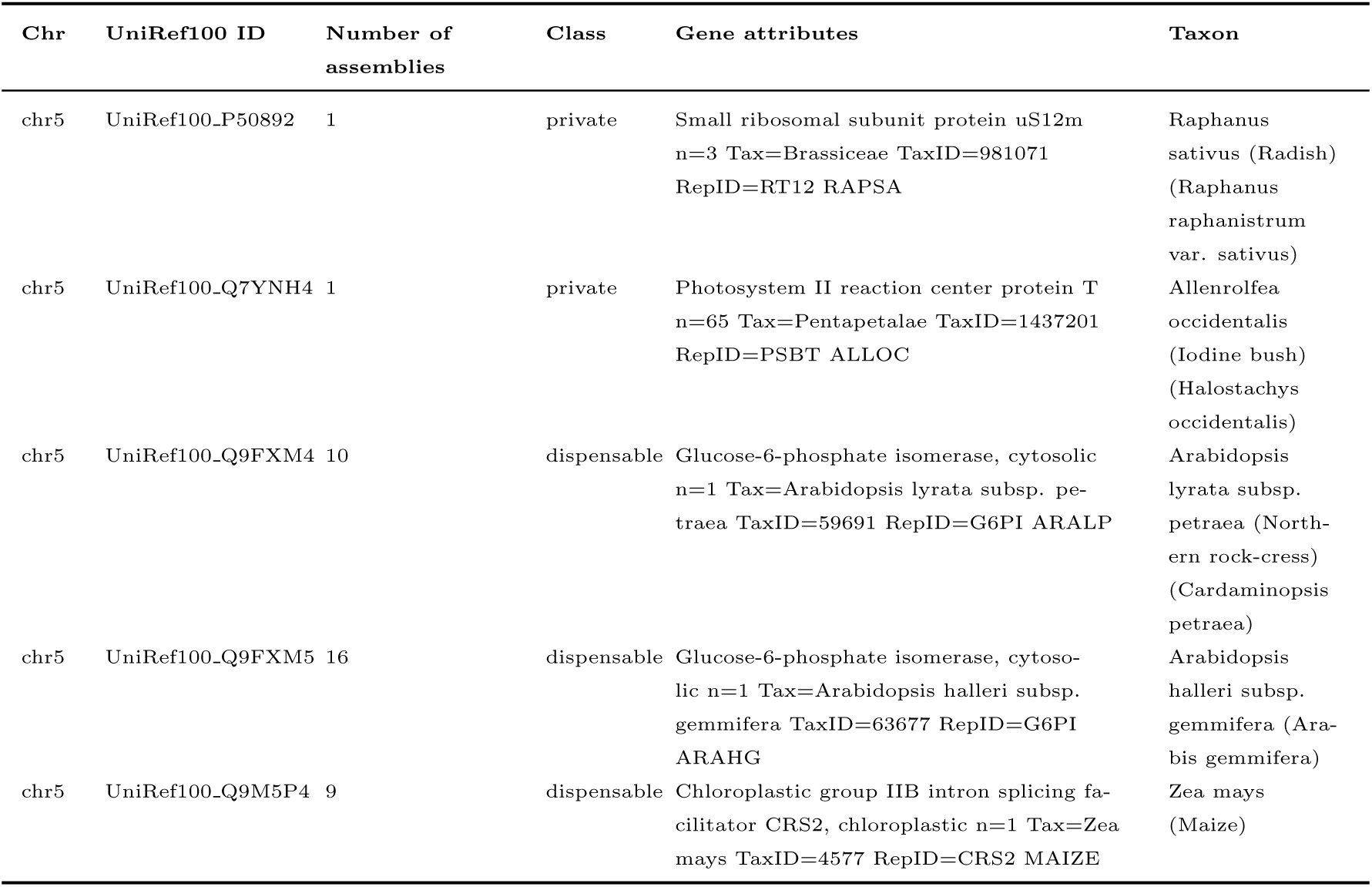
New pseudogenes found in non-reference sequences attributed to species different from *A. thaliana*.

One of the new pseudogenes, originating from the parent gene UniRef100 Q75VR0 and the species *Nicotiana tabacum*, was categorised as softcore and found in 80 assemblies. 28 new pseudogenes originating from species different from *A. thaliana* were classified as dispensable, and 13 as private.

The pseudogene UniRef100 Q332S9 from *Lactuca sativa* was found in a total of 5 accessions, but on different chromosomes (chromosome 3 for GCA 933208065, GCA 946403525, GCA 946408975; and chromosome 4 for GCA 946403025, GCA 946412065), and therefore was counted as a different pseudogene twice.

We explored the gene ontology enrichment of the pseudogenes classified into core and variable (i.e., softcore, dispensable, and private) using TopGO.

The gene ontology of pseudogenes was analysed by hypothesizing that the parent genes are the best approximation of pseudogene function at the time of their origin.

Thus, based on parent gene ontology, variable (softcore, dispensable, and private) pseudogenes were significantly enriched for terms related to a wide range of processes, including not only primary developmental processes such as morphogenesis, reproductive development, and regulation of vegetative growth but also response to abiotic stresses and to xenobiotic stimulus. We found no significant enrichments for the GO terms of core pseudogenes. (Supplementary Materials Figure S4).

The similarity tree obtained considering the Jaccard distance between accessions based on pseudogenes PAVs is displayed in Figure 15. The tree’s tips were coloured according to the country of origin of the accessions to highlight potential geographic-specific clusters. The tree’s robustness was confirmed by a cophenetic correlation coefficient equal to 0.86. The Mantel test yielded a correlation coefficient of 0.5469028 and a simulated p-value of 0.0001, indicating a strong positive and statistically significant correlation between the genetic distance in terms of pseudogenes and the geographic distance between the accessions.

**Figure 15:**
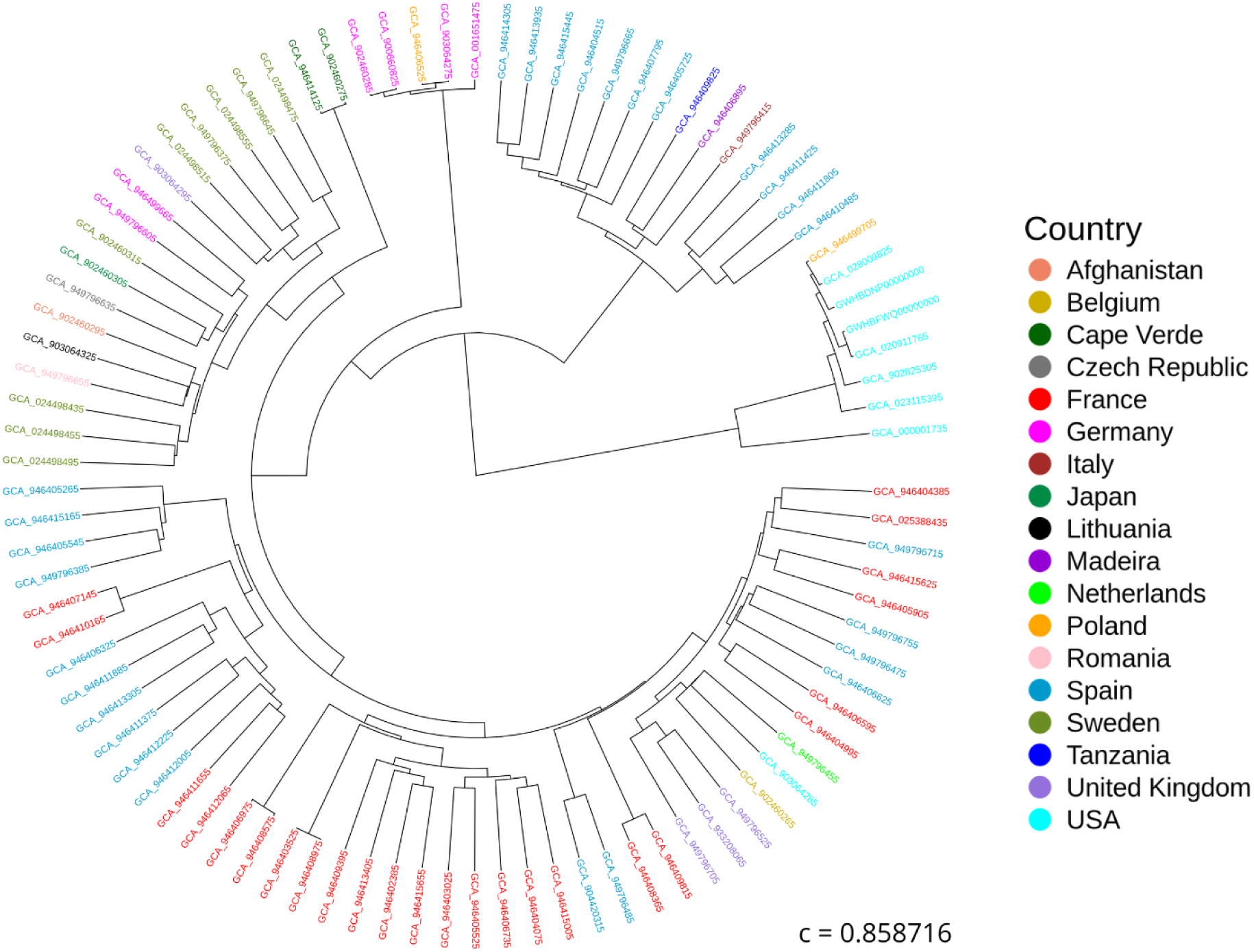
Similarity tree based on the Jaccard distance between assemblies calculated on the pseudogene PAVs. “c” represents the cophenetic correlation coefficient.

## 4 Discussion

This study describes the first *A. thaliana* pangenome graph constructed with a reference-free approach PGGB [23]. Moreover, it included 93 assemblies, representing a significative size improvement over previous pangenomes.

Reference-based methodologies may yield biased outcomes that reflect an incomplete representation of genetic variation [54].

Jiao et al. (2020) [18], which constructed the pangenome using 8 accessions and applying a pairwise whole-genome sequence alignments of all possible pairs of genomes method, estimated the size of the *A. thaliana* pangenome to be 135 Mb. Kang et al. (2023) [19] conducted the sole prior study employing a graph-based methodology to develop a pangenome of this species. They employed the Minigraph method [21], which uses a reference genome as backbone to construct the graph, using 32 assemblies. The pangenome size was 243.27 Mb, comprising 468,168 nodes and 649,692 edges. Our pangenome included over double the sequence length (488.78 Mb), comprising 30,391,243 nodes and 7,189,634 edges, thereby capturing a greater degree of variation. This outcome is certainly a result of the higher number of assemblies included, but it is also the result of the method employed, which capture a higher degree of diversity.

A total of 31,017 genes and 23,894 pseudogenes were successfully mapped to the pangenome graph. The overall variability observed among the assemblies was remarkable: 65% of nodes, 30% of genes, and 95% of pseudogenes were categorised as either dispensable or private.

Kang et al. (2023) [19] annotated a total of 887,723 genes across 32 assemblies, resulting in an average of 27,741.3 genes per assembly. Lian et al. (2024) [20] annotated a total of 1,928,005 across 69 assemblies, averaging 27,942.1 genes per assembly. With our approach, we were able to map a total of 2,325,577 genes across 93 assemblies, with an average of 25,006.2 genes per assembly. Although our average yield is lower compared to the previous studies, it should be appreciated that it does not require previously annotated genomic sequences. Furthermore, we could map a higher number of genes if we applied a less strict filter to the results derived from mapping the reference genes to the pangenome graph, which was 95% identity and 95% coverage. Overall, annotating the pangenome graph allows to obtain a global view of the variation within the species and to study intra-specific diversity, requiring lower efforts compared to the approach applied in previous studies.

Previous pangenomic studies on other plant species employed methods that are not directly comparable to our approach. The employment of the reference-free approach used in the current study could explain the substantial increase in size of the pangenome over the reference genome — 310.83% larger than the reference genome — compared to previous studies. For example, the pangenome of *Oryza sativa*, which was constructed from 111 accessions using a long-read sequencing-based approach, exhibited an increase of 156% over the reference genome [55]. Similarly, other studies reported increases of 0.81% in *Glycine soja* [56], 20% in *Brassica oleracea* [57], 3.3% in *Triticum aestivum* [58], 22% in *Brassica napus* [59], and 24% in *Sorghum bicolor* [60].

Regarding the size of genomic classes, previous studies on different plant species showed significant variability in the proportion of core genes. For instance, core genes accounted for 48.6% of the total in *Glycine soja* [56], 81.3% in *Brassica oleracea* [57], 64.3% in *Triticum aestivum* [58], 47% in *Sorghum bicolor* [60], 65.2% in *Brassica napus* [59], 89.5% in *Amborella trichopoda* [61], 58.21% in *Sesamum indicum* [50], 70% in *Cucumis sativus* [62], 19.01% [63] and 30% [64] in *Oryza sativa*, and 36% in *Glicine max* [65]. These results underscore the extensive variability across species. Furthermore, most of these studies classified gene PAVs based on gene family clustering rather specific loci, as done in this study. This underscores the importance of the method employed in our study, which allows for a more precise classification of features into genomic classes.

We found that the variation observed among *A. thaliana* accessions was partially correlated to the geographical distance among the locations where the samples were collected. As regards to the node-based dissimilarity, we found that there was no correlation with the difference in geographical origin, suggesting that genome structure is overall conserved, as it was previously found by Lian et al. (2024) [20]. However, we observed that the diversity among accessions was correlated to the distance among their geographical origins in terms of genes and especially pseudogenes. Sisu et al. (2014) [66] pointed out that the majority of pseudogenes evolve neutrally because, unlike functional genes that are essential for the development and functioning of an organism, they are not under selective pressure. This makes pseudogenes an ideal resource to study the impact of demographic forces on genome evolution, with potential applications in future pangenomic studies.

We observed the presence of mitochondrial and chloroplast genes in the pangenome, even though assembled organelles were initially excluded from the assemblies. This result could be attributed to errors in the assembly process caused by their incorrect removal from nuclear sequences. However, some of these may represent real insertions. For example, the largest number of mitochondrial genes was found in chromosome 2, where a large insertion of mitochondrial DNA was previously documented [67]. Insertions from both mitochondrial and chloroplast DNA in the nuclear genome were observed on all chromosomes of *A. thaliana* in previous studies [68, 69].

The simulation curve for pangenome size growth (Figure 11) revealed that we had nearly reached a plateau with 93 assemblies. Kang et al. (2023) [19] previously showed that they approached a plateau with 32 assemblies; however, Lian et al. (2024) [20]affirmed that they could not capture all the diversity with 73 assemblies since their pangenome growth curve had not yet reached a plateau. Despite our curve seeming to be close to a saturation point, further diversity might still be found by adding new accessions to the pangenome.

## 5 Conclusions

In this study we presented a reference-free pangenome of the model plant *A. thaliana*. We invested significant effort in developing methods to analyse the direct mapping of genes and pseudogenes to the pangenome graph, and to find a way to annotate the sequences that were not shared with the reference genome. The assemblies we used were not produced in this study, and therefore we could not be completely sure about their quality. The overall methods still need improvement, but our findings show the potential of reference-free pangenomes in uncovering intra-specific diversity, even in a species as well-known as *A. thaliana*. We believe that this study contributed to our comprehension of intra-specific variation in *A. thaliana* and will offer a novel perspective on the potential applications of pangenomes in comparative genomics.

## Data and code availability

The genome assemblies used in this study are available at the NCBI and CNCB databases under the project IDs reported in Supplementary Materials Table S1. The scripts developed for this study and the command used are available on GitHub at https://github.com/LiaOb21/arabidopsis_pangenome.

## Authors’ contributions

LO performed the bioinformatic analyses described in this work, designed the Phyton scripts, and wrote the first version of the manuscript. AG provided substantial support for the pangenomics analysis. AP designed the Perl scripts used in this work and performed the Gene Ontology enrichment analysis. UT and AP provided supervision and guidance throughout the project. LO, UT, and AP contributed to the manuscript.

## Funding

This study includes part of a PhD project carried out by Lia Obinu at the PhD School of Agricultural Sciences of the University of Sassari, funded by the University of Sassari. Lia Obinu was also supported by the ERASMUS for Traineeship program awarded to the University of Sassari. This study was carried out within the Agritech National Research Centre and received funding from the European Union Next-Generation EU (PIANO NAZIONALE DI RIPRESA E RESILIENZA (PNRR) - MISSIONE 4 COMPONENTE 2, INVESTIMENTO 1.4 - D.D. 1032 17/06/2022, CN00000022). This manuscript reflects only the authors’ views and opinions and neither the European Union nor the European Commission can be considered responsible for them.

## Notes

### Competing Interest Statement

The authors have declared no competing interest.

