## Supplementary material for "The reference-free pangenome of *Arabidopsis thaliana*": Figure_S1_Arabidopsis_threshols_trial_chr1.pdf

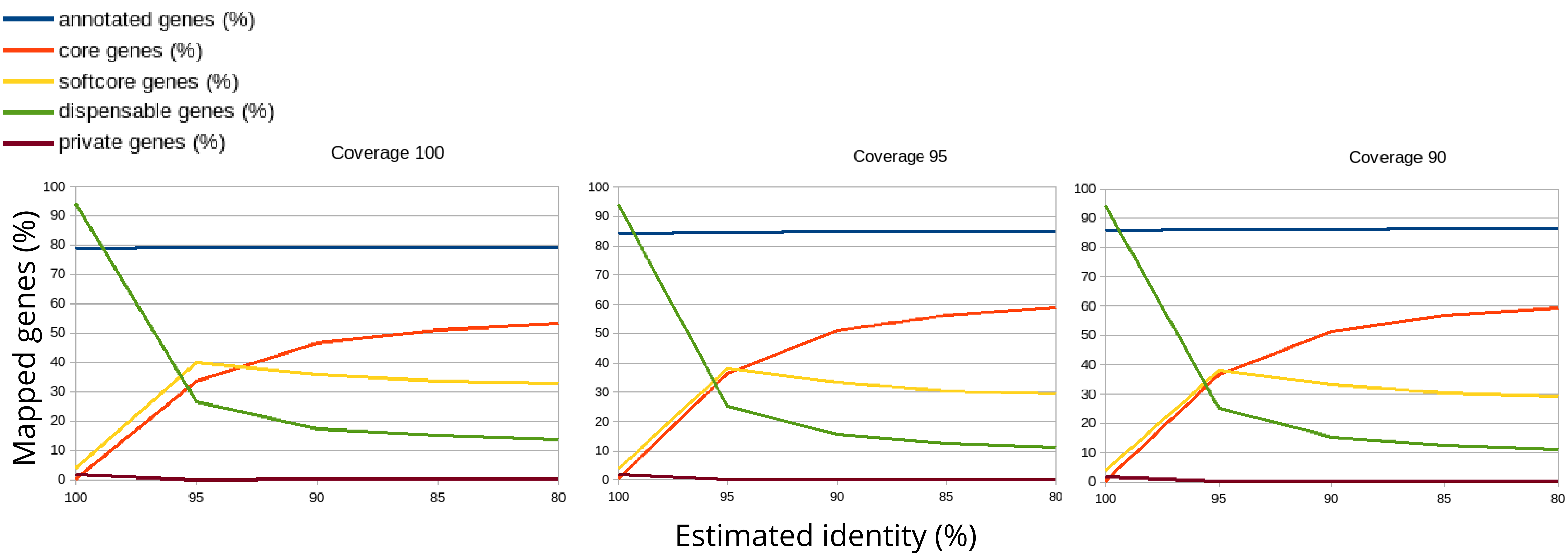


Figure S1. Percentage of genes mapped to the community corresponding to chromosome 1 applying different thresholds for coverage and estimated identity in A. thaliana pangenome.
