## Supplementary material for "The reference-free pangenome of *Arabidopsis thaliana*": Figure_S3_TopGO_variable_genes.pdf

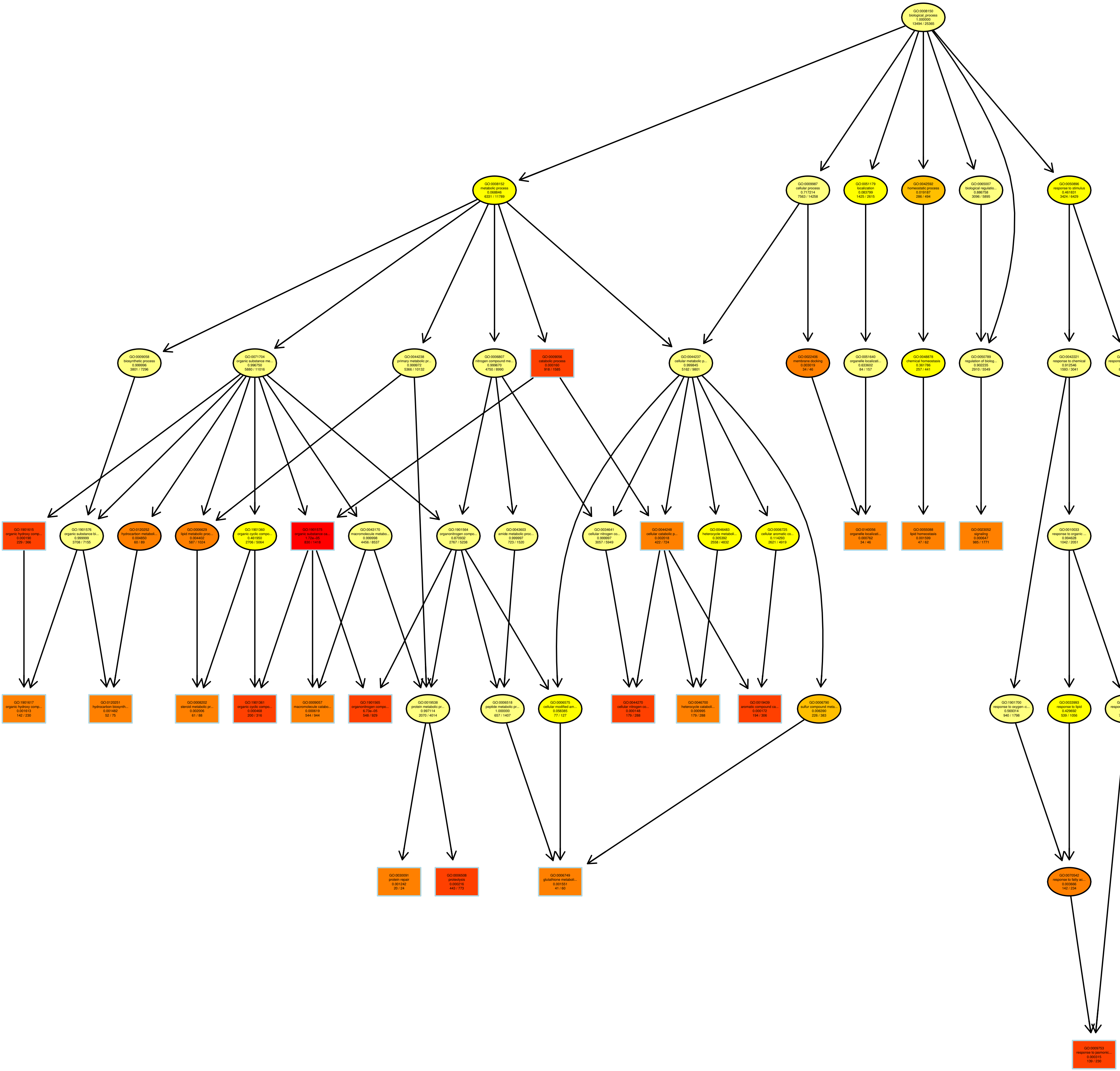

*Figure S3. GO enrichment analysis of variable genes(i.e., softocore, dispensable, and private) in the A.thaliana pangenome.*
