## Supplementary material for "The reference-free pangenome of *Arabidopsis thaliana*": Figure_S4_TopGO_variable_pseudogenes.pdf

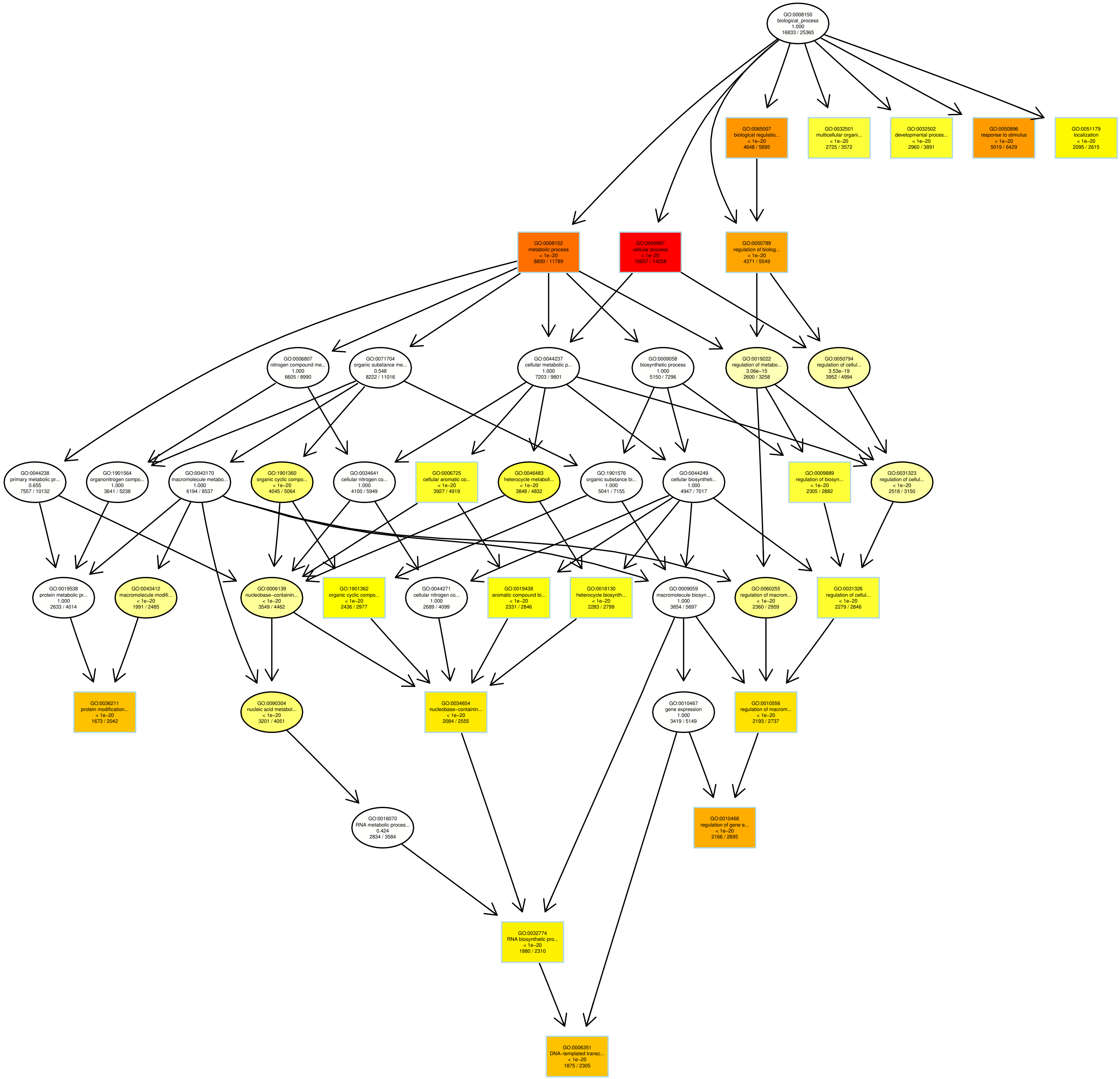


*Figure S4. GO enrichment analysis of variable pseudogenes (i.e., softocore, dispensable, and private) genes in the A.thaliana pangenome.*
