## Supplementary figures and images for "The reference-free pangenome of *Arabidopsis thaliana*"

### Figure_S2_TopGO_core_genes.pdf

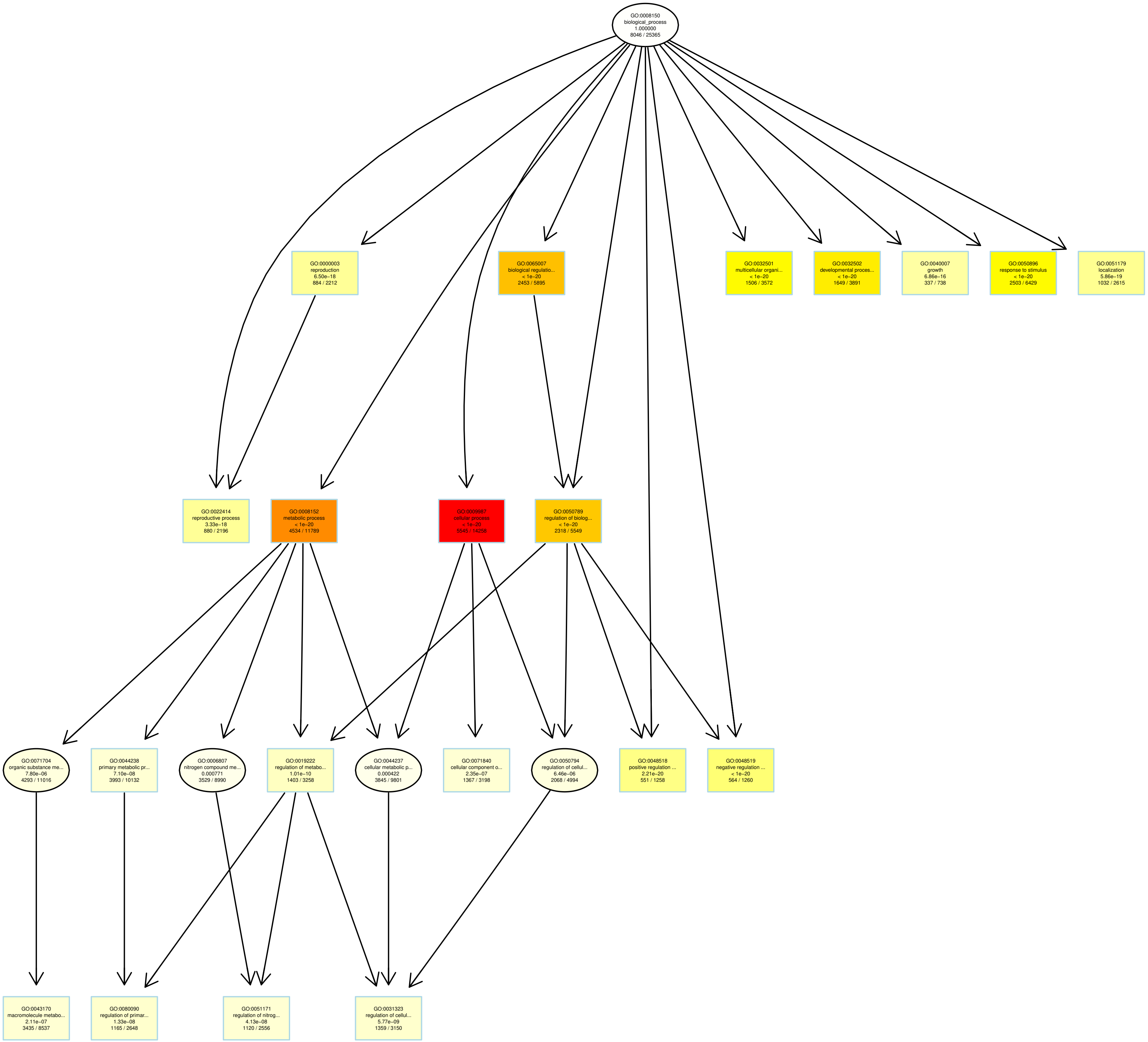


*Figure S2. GO enrichment analysis of core genes in the A. thaliana pangenome*
